# Next-Generation Nanopore Sensors for Enhanced Detection of Nanoparticles

**DOI:** 10.1101/2023.03.31.534385

**Authors:** Samuel Confederat, Seungheon Lee, Der Vang, Dimitrios Soulias, Fabio Marcuccio, Timotheus I. Peace, Martin Andrew Edwards, Pietro Strobbia, Devleena Samanta, Christoph Wälti, Paolo Actis

## Abstract

Nanopore sensing is a technique based on the Coulter principle to analyze and characterize nanoscale materials with single entity resolution. However, its use in nanoparticle characterization has been constrained by the need to tailor the nanopore aperture size to the size of the analyte, precluding the analysis of heterogenous samples. Additionally, nanopore sensors often require the use of high salt concentrations to improve the signal-to-noise ratio, which further limits their ability to study a wide range of nanoparticles that are unstable at high ionic strength.

Here, we report the development of nanopore sensors enhanced by a polymer electrolyte system, enabling the analysis of heterogenous nanoparticle mixtures at low ionic strength. We present a finite element model to explain the anomalous conductive/resistive pulse signals observed and compare these results with experiments. Furthermore, we demonstrate the wide applicability of the method by characterizing metallic nanospheres of varied sizes, plasmonic nanostars with various degrees of branching, and protein-based spherical nucleic acids with different oligonucleotide loadings.

Our system will complement the toolbox of nanomaterials characterization techniques and will enable real-time optimization workflow for engineering a wide range of nanomaterials.

## INTRODUCTION

Over the past few decades, the use of nanoparticles has played a significant role in the advancement of medicine, optics, and electronics (1-3). The use of nanoparticles not only sparked a strong engagement in the research settings, but they have also become widely incorporated in numerous consumer goods nowadays (4). Understanding the structural– functional relationships of engineered nanoparticles is a continuous undertaking that requires an in-depth exploration of their physicochemical properties. Therefore, the ability to characterize nanoparticles in a high-throughput manner is of utmost importance. However, characterizing nanoparticles in their native state, specifically in heterogenous mixtures, presents many challenges (5). Dynamic light scattering (DLS) or UV-Vis spectroscopy are ensemble-averaging techniques and, therefore, fall short in fully characterizing heterogenous nanoparticle mixtures (6). Nanoparticle tracking analysis (NTA) is suitable for analysing size distribution of polydisperse nanoparticle suspensions with single entity resolution, however the nanoparticles need to have a reflective index distinct from the surrounding medium or a fluorescent label is required (7). Imaging methods such as transmission electron microscopy (TEM) provide high-resolution characterization of individual nanoparticles but suffer from sampling bias, low throughput, and require careful sample preparation (8).

Nanopore sensing is a powerful label-free electrical technique that uses the Coulter principle for single entity analysis (5). In nanopore experiments, individual entities are driven through a nanopore under the influence of an electric field, causing a temporary modulation in the recorded ion current by a combination of geometrical exclusion of the electrolyte solution, ion concentration polarization, and additional charges brought by the analyte itself (9,10). The magnitude and duration of these modulations reflect the translocation dynamics of the analyte, which are dependent on its physicochemical properties (e.g., size, shape, charge) (11-13). Even though nanopores have been employed in numerous sensing applications, nanopore technology has an untapped potential for the analysis of nanoparticles (14). This is because current nanopore measurements require the size of the pore to match the size of the analyte, limiting the investigation to homogenous mixtures (15-17). Furthermore, nanopore measurements often require high-ionic-strength electrolytes which precludes the analysis of nanoparticles systems that are unstable at high ionic strength (18,19). Tuneable resistive pulse sensing (TRPS) is a nanopore technique that has found applications in nanoparticle characterization, but it is limited to nanoparticles larger than 100 nm in size (20,21). Furthermore, the TRPS signals deviate from linearity when the size of the nanoparticle approaches the diameter of the pore aperture, limiting its analytical capabilities.

Here, we present polymer-electrolyte enhanced nanopore sensing which enables the analysis of heterogenous nanoparticle samples at low ionic strength. The polymer electrolyte environment generates a large signal enhancement eliminating the need for a nanopore that matches the size of the nanoparticle and therefore allowing the high throughput analysis of heterogenous nanoparticle mixtures. Furthermore, combining experimental findings and finite-element modelling (FEM), we provide a mechanistic explanation for the ion current signatures. We demonstrate the characterization of nanoparticles at low ionic strength (10 mM KCl) enabling the analysis of anisotropic gold nanostars (AuNS) with varying degrees of branching and report the detection and analysis of an emerging class of functional soft nanoparticles, protein spherical nucleic acids (ProSNAs) with distinct oligonucleotides shells. The single-nanoparticle analysis approach described herein will complement the toolbox of existing nanomaterials characterization techniques, unleashing the potential of nanopore sensing for the universal analysis of nanoparticles.

## RESULTS AND DISCUSSION

A polymer electrolyte nanopore system (22) enables the detection of heterogenous nanoparticle mixtures with a fixed nanopore size (23). We fabricated glass nanopores with a diameter of 60 nm and probed the translocation of 20 nm diameter gold spherical nanoparticles (AuNPs) samples (Figure 1) under an applied voltage of −500 mV. The nanopore setup consisted of a glass nanopore filled with and immersed in an electrolyte (KCl) solution where the application of a potential between a pair of Ag/AgCl electrodes, inside the glass pore and external bath respectively, drive the translocation of the analyte towards the external bath. As shown in Figure 1A, no translocation events were observed for the 20 nm AuNPs using standard electrolyte buffer conditions (i.e., 50 mM KCl solution). However, the addition of 50% w/v PEG (polymer electrolyte) to the outer bath resulted in conductive translocation events well-resolved from the ion current baseline (Figure 1B). These events are characterised by a specific peak amplitude (ΔI_c_) and dwell time (Figure 1B inset), where each peak is generated by individual nanoparticles translocating from inside the pore to the outer bath. No events were observed under the application of a positive voltage or in absence of nanoparticles in solution (Figure S5). Increasing the magnitude of the applied voltage led to an increase in amplitude of the conductive peak current and a decrease in the dwell time, suggesting that the electrophoretic force plays a major contribution in driving the negatively charged AuNPs through the pore (Figure S6). Similarly, we observed a linear increase in the frequency of the translocation events with increasing voltage and increasing nanoparticle concentrations (Figure S6).

**FIGURE 1.**
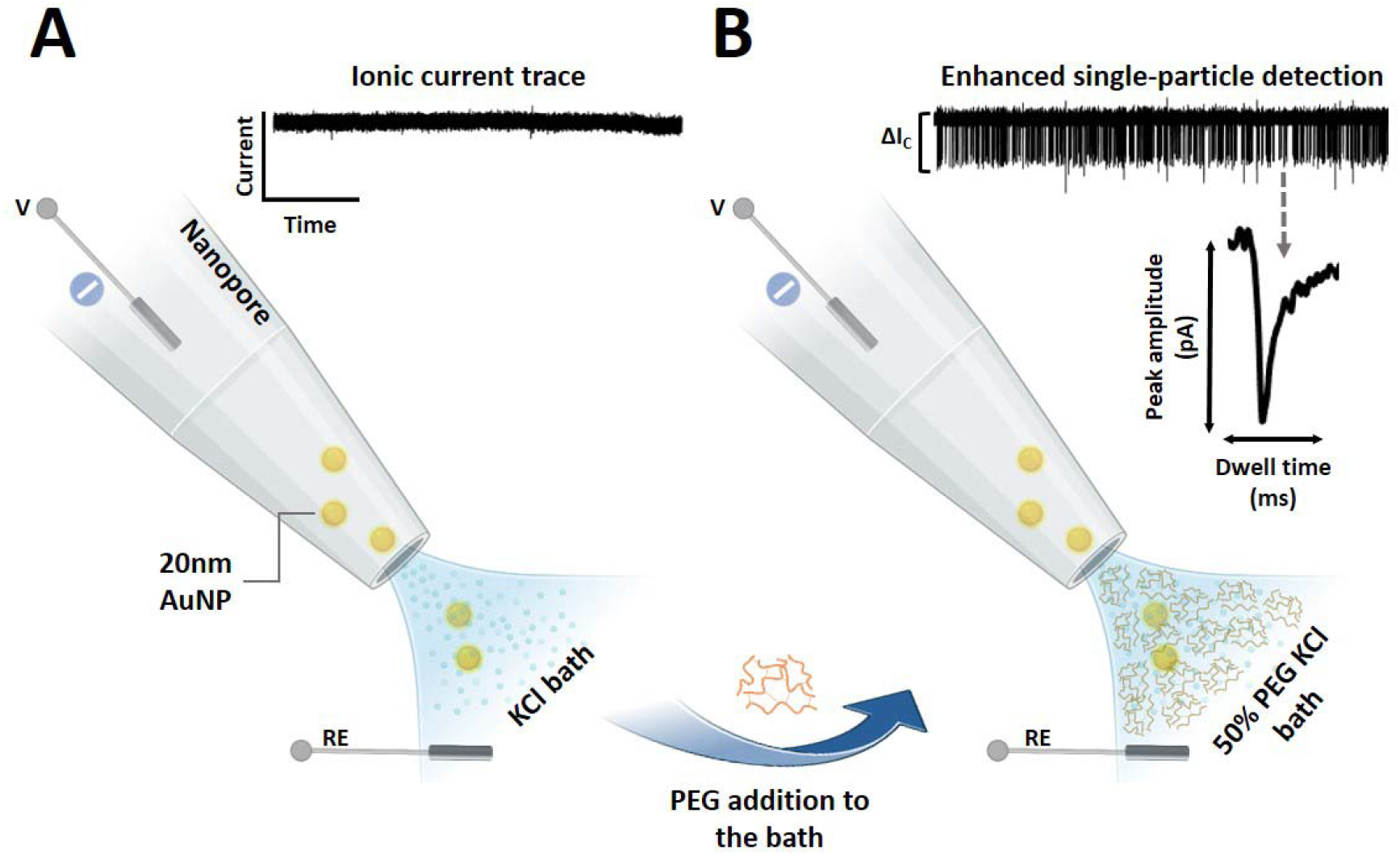
Enhanced nanoparticle detection in polymer electrolyte bath **(A)** Schematic illustration depicting 20 nm diameter AuNPs translocating across the nanopore towards the outer bath into the outside and the recorded ion current trace at −500 mV for 20 nm AuNPs utilizing a 50 mM KCl external electrolyte bath. No translocation events were observed under these conditions. **(B)** Representative translocation setup depicting the use of 50% (w/v) PEG (polymer electrolyte) in 50 mM KCl in the external electrolyte bath with the corresponding recorded ion current trace at −500 mV, showing conductive translocation events for 20 nm AuNPs. A representative translocation event is shown with the translocation peak amplitude and dwell time characteristics indicated.

Figure S3 depicts the translocation of a 50 nm diameter and 20 nm diameter AuNP mixture where two distinct peak amplitudes can be observed that also result in distinct populations in the events scatter plots (Figure S3C). The average peak amplitude was 61.1 ± 1 pA for the 20 nm AuNPs and 225 ± 1 pA for the 50 nm AuNPs. The large amplitude difference denotes the strong signal enhancement generated by the polymer electrolyte that allowed us to further probe a range of different AuNPs (50, 40, 30, 20, and 10 nm diameter) with a fixed nanopore size (60 nm in diameter) as shown in Figure 2A. In contrast to previous nanopore strategies employing chemical modifications (24) or arrayed nanopores (25), our nanopore system supports rapid detection of heterogenous nanoparticles and clear discrimination of their diameter size with a fixed pore diameter size. Whereas previous studies demonstrated the discrimination of nanoparticle mixtures utilizing a single nanopore, they were predominantly used for the analysis of binary mixtures with relatively large difference in their size (26,27). Here, we further explored the discrimination of nanoparticles with 10 nm size difference in the low nanometre range (10 – 50 nm). The translocation signal distributions were fitted to Gaussian curves which enabled the identification of 5 distinct populations, one for each nanoparticle set (Figure 2B). The average conductive peak current increased with the ratio of nanoparticle to pore diameter (d_Np_/d_pore_), as shown in Figure 2C. To further support the capability of our nanopore approach to discriminate heterogeneous nanoparticles mixtures, in Figure S7 we compared the translocation of individual AuNP solutions with a mixture containing all AuNPs (50, 40, 30, 20, and 10 nm) in one solution.

**FIGURE 2.**
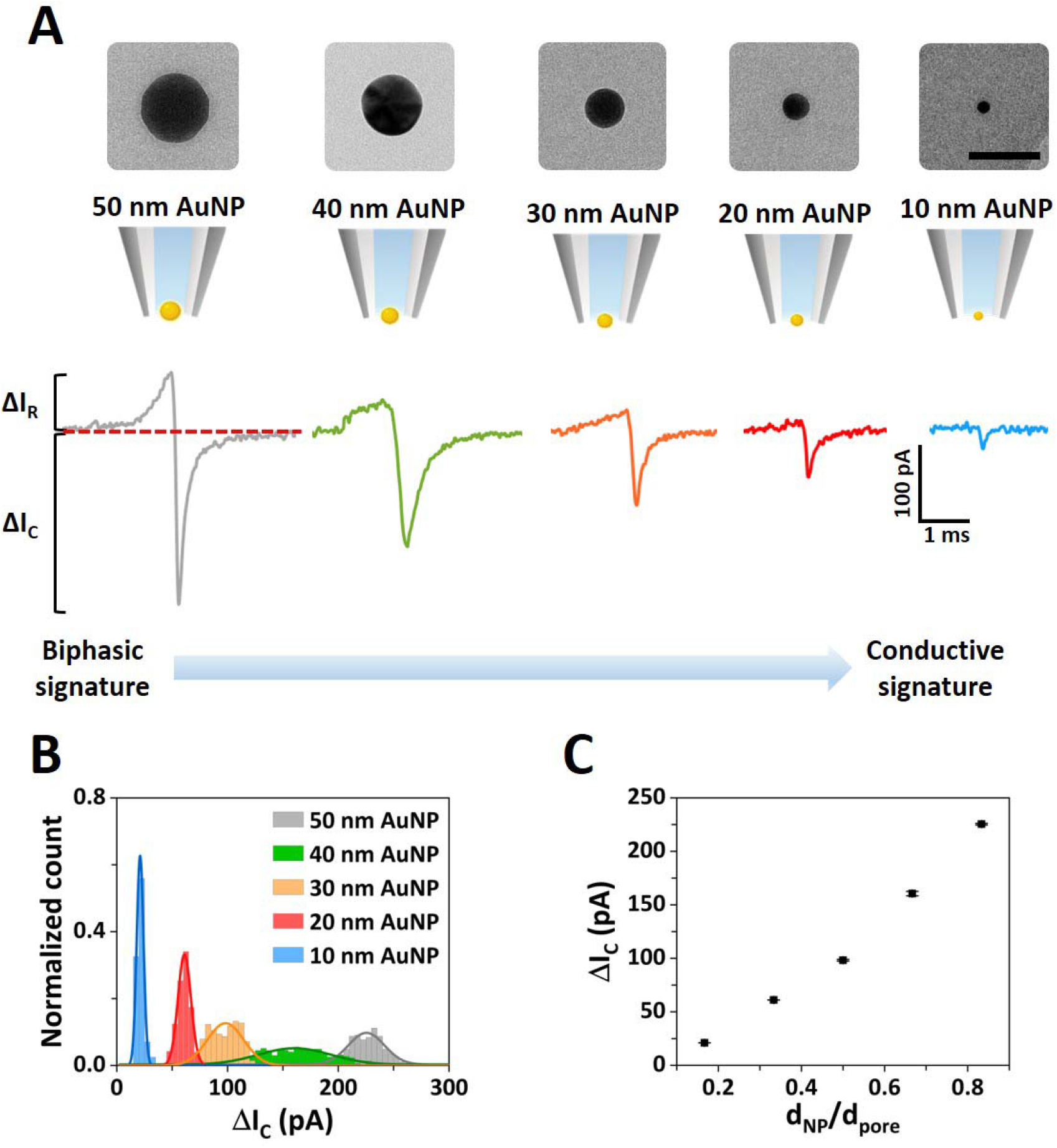
Nanopore detection of gold nanoparticle mixtures. **(A)** Top panel: TEM images of the standard AuNPs, left-to-right: 50, 40, 30, 20, and 10 nm diameter (50 nm scale bar). Bottom panel: schematic representation of the AuNP translocating through a 60 nm diameter fixed pore size with representative individual translocation peaks for each AuNP, denoting a transition from biphasic to conductive signature when the ratio of nanoparticle to pore size decreases. **(B)** Event histogram of the conductive peak current for the five different AuNPs (50, 40, 30, 20, and 10 nm diameter) translocated with a fixed pore size (∼60 nm diameter). The solid lines represent Gaussian fits to each translocation data set. **(C)** Average conductive peak current as a function d_Np_/d_pore_. The error bars represent the standard error.

Whereas the conductive peak represents the dominant feature in the ion current signatures, we observed a distinct change in the translocation signature when different nanoparticle to pore size ratios were employed. Namely, when the size of the nanoparticle closely matched the size of the pore, an additional resistive peak was observed (Figure S8). This behaviour is evidenced in Figure 2A, where a biphasic (resistive and conductive peak) signal transitions to a conductive signal when the size of the AuNP decreases with respect to the size of the pore (Table S1) or vice versa (Figure S9). Classically, nanoparticle analysis with nanopores leads to resistive (current-decreasing) translocation signals because of the ion flow hindrance by the presence of the nanoparticle within the nanopore sensing region (26). However, several studies have shown the occurrence of conductive peaks. Sensale et al. identified the surface charge of the particle as the main factor that influences the characteristics of the translocation signal through a glass nanopores (28). The authors suggested that this phenomenon occurs due to an ion accumulation/depletion associated with the surface charge of the particle translocating through the pore. Similarly, in the studies conducted by Menestrina et al. and Chen et al., biphasic signals were reported both experimentally and in simulations when charged particles are translocated in low salt buffer conditions (below 200 mM KCl) (29,30). The authors reported resistive peaks followed by a conductive component and explained the effect considering the charge carried by the analyte that causes a temporary ion enrichment at the aperture when low-salt conditions are used. The phenomenon of the current enhancement can be influenced by several factors, including the electrolyte concentration, nanoparticle to pore size ratio, and the nanoparticle surface charge (31). To investigate the origin of the conductive contribution to the peak in our nanopore system, we developed a finite-element model to estimate the potential resulting current enhancement when a polymer electrolyte (50% PEG) is used in the external solution (32). We hypothesised that an interface is formed at the nanopore tip between the inner and outer solution (KCl only and the polymer electrolyte bath), and that this interface is deformed by a model nanoparticle (50 nm diameter) translocating through the pore. This deformation induces an ionic rearrangement within the nanopore producing the single nanoparticle events enhancement. Interestingly and counter-intuitively the presence of the nanoparticle in the nanopore sensing area leads to an increase in the ionic concentration due to the deformation of the interface which far outweighs the volume-exclusion-related decrease, thereby overall yielding a temporary current increase which manifests in a conductive peak. To further highlight the influence of the interface on the conductive peak enhancement, we simulated the effect of a negatively charged nanoparticle versus a neutral nanoparticle of similar size. As shown in Figure S19, the magnitude of the conductive peak current is largely influenced by the external polymer electrolyte interface, with small contribution exerted by the particle surface charge. Remarkably, although we simulated the effect of the presence of a nanoparticle at distinct locations within the nanopore at equilibrium for each position (Figure 3A), we observed an excellent agreement with the experimentally measured resistive and conductive signal (Figures 3B and 3C). We note that we do not have experimental data on the nature of the interface at the nanopore between the inner and the outer bath and therefore assumed it to be a straight boundary at z = 0 nm. This is clearly an over-simplification, and the interface geometry is likely to be more complex. Also, our stationary and ergodic model may not capture the full dynamics of the nanoparticle translocation across the interface, but it does capture the underlying process and provides a good phenomenological understanding of the impact of the nanoparticle translocation on the ion current. Using a more sophisticated fully dynamic model would certainly improve the accuracy of the predictions but would unlikely lead to a significant revision of the phenomenological understanding of the system.

**FIGURE 3.**
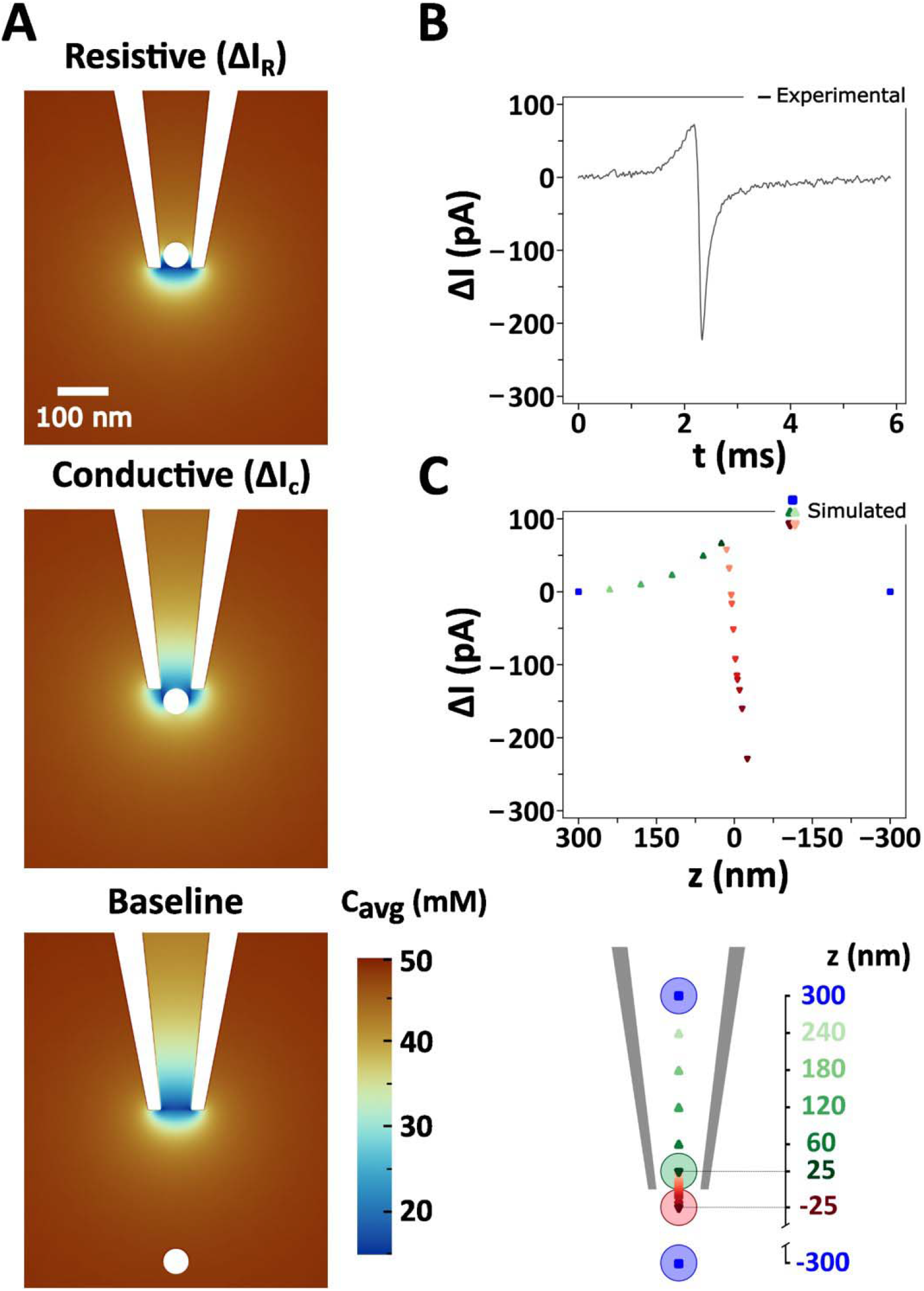
Finite element simulations of AuNP translocation. **(A)** Simulated ion distribution in the proximity of the nanopore tip utilizing a 50% PEG in 50 mM KCl electrolyte bath with a 50 nm diameter AuNP at different translocation positions through a 60 nm diameter pore (top: inside the nanopore, middle: at the nanopore tip, bottom: outside in the bath). Experimental **(B)** and simulated **(C)** translocation signal of the 50 nm diameter AuNP through a 60-diameter nm pore with 50% PEG in 50 mM KCl electrolyte bath, and a nanopore schematic depicting the translocation distance used for simulated translocation signal (bottom panel).

Our polymer electrolyte system also enables the analysis of nanoparticles at low ionic strength (10 mM). Generally, nanopore measurements are carried out in high-salt conditions (> 100 mM KCl), excluding the analysis of less stable nanoparticles, such as citrate-capped nanoparticles (33). We probed the translocation of gold and platinum citrate-capped nanoparticles and bare silver nanoparticles diluted in 10 mM KCl, demonstrating that our nanopore measurements can reliably detect nanoparticles with high-capture rates and high signal-to-noise ratios (SNR). These results are evident in Figure 4A, where the translocation of three sets of 30 nm metallic nanospheres (Figure S10) utilizing a 60 nm nanopore biased at −500 mV. The data also shows a good agreement between the average conductive peak current recorded for each set of nanoparticles with nominally the same size (30 nm diameter). The Gaussian fits of the peak current distribution in Figure 4A resulted in average conductive peak current of 40 ± 1 pA for the AuNPs, 40 ± 1 pA for the PtNPs, and 38 ± 1 for the AgNPs, respectively.

**FIGURE 4.**
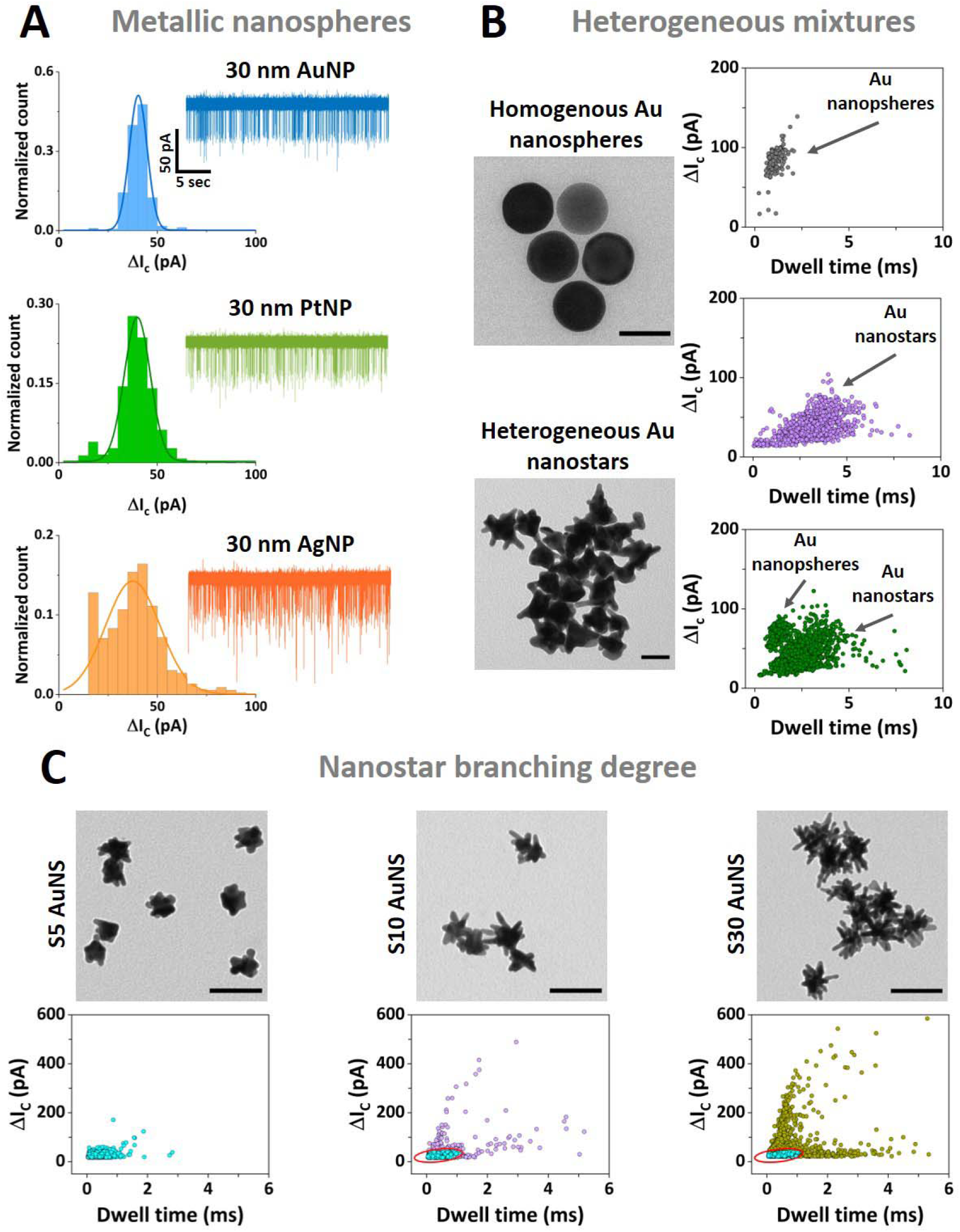
**(A)** Nanopore sensing of metallic nanospheres with histograms of the conductive peak current distribution and representative ion current traces for: 30 nm diameter PEG carboxyl-capped gold nanospheres (top panel), 30 nm diameter citrate-capped platinum nanospheres (middle panel), and 30 nm diameter citrate-capped silver nanospheres (bottom panel). The solid lines represent Gaussian fits to each translocation data set. Nanopore recordings were carried out in 50% PEG in 10 mM KCl utilizing a 60 nm diameter fixed pore size biased at −500 mV. The current and time scales are the same for all the ion current traces. **(B)** Nanopore sensing of homogenous gold nanospheres (TEM top panel) and heterogenous gold nanostars (TEM bottom panel). Scale bar TEM graphs: 50 nm. Scatter plots with conductive peak current (ΔI_C_) versus dwell time of the translocation events are depicted for the individual sets (top panel: gold nanospheres; middle panel: gold nanostars), and a mixture (bottom panel: gold nanospheres and gold nanostars) in similar nanopore conditions (50% PEG in 10 mM KCl using an 80 nm diameter fixed pore biased at −500 mV). **(C)** Nanopore sensing of nanostars with different branching degree of three sets of synthesized gold nanostars as depicted in the TEM images: S5 (left panel), S10 (middle panel), and S30 (right panel) AuNS (100 nm scale bar). The notation S5-S30 denotes their different branching degree (from low to high). Scatter plots of conductive peak currents as a function of dwell time for the S5 AuNS sample (left panel), S10 AuNS sample (middle panel), and S30 AuNS sample (right panel). The red ellipse in the S10 and S30 scatter plot data indicates the 95% confidence interval of the S5 translocation events. 12% of the events fall outside the S5 CI for the S10 sample and 35% of the events fall outside the S5 CI. Nanopore recordings carried out in 50% PEG in 10 mM KCl conditions utilizing 80 nm diameter pores biased at −700 mV.

Inspired by the results above, we expanded our nanopore measurements to tackle complex anisotropic nanoparticles, such as branched gold nanostars (AuNS). AuNS are emerging as a prominent plasmonic particle for application in surface-enhanced Raman scattering (SERS) and offer several advantages over spherical nanoparticles (34-36). The localized surface plasmon in AuNS is tuneable by controlling the anisotropy of the structure during the synthesis (37). However, current characterization and quality control for these plasmonic nanostructures relies mainly on TEM imaging. We first probed a nanoparticle solution composed of AuNS spiked with 50 nm gold nanoparticles. Their nominal hydrodynamic radius was broadly similar (60 nm vs 50 nm), but the ion current signals generated two clearly distinct populations indicating the potential of our approach for the quantitative analysis of anisotropic nanoparticles (scatter plots Figure 4B). Opposite to the translocation events obtained for the uniform gold nanospheres, the events recorded for the AuNS samples show a wider spread in terms of the peak current and dwell time. We attribute these differences to the irregular shape of the synthesized nanoparticles and their heterogenous character, as depicted in the TEM images in Figure 4B. Furthermore, we probed suspensions of AuNS with low and high degree of branching in 50% PEG and 10 mM KCl following their different synthesis stages (Figure 4C). Similarly, we use a fixed nanopore size (80 nm diameter) to probe the translocation of distinct AuNS samples (Figure 4C), here named S5, S10, and S30, respectively (see gold nanostar synthesis section), according to their degree of branching. The samples all exhibit a large amount of anisotropy which gives rise to their unique optical properties (Figure S12 and Figure S13). With increasing branching density, we observed a broadening of the distributions both in terms of peak current amplitude and dwell time as shown in the scatter plots in Figure 4C and Figure S11. To evidence the progression in terms of nanopore detection from synthesis of AuNS (S5) to the high-density branched AuNS (S30) we computed a 95% confidence interval (CI) using the S5 translocation events as an input. We then applied this CI to the translocation events obtained for S10 AuNS and S30 AuNS samples. Based on this CI fitting we outlined an increase in the percentage (from the total number of events) of the events falling outside the S5 CI, namely 12% for the S10 and 35% for the S30 AuNS. In future, more sophisticated analysis employing algorithmic models could reveal more subtle information concerning the ion current signals.

To further expand on the applicability of our polymer electrolyte enhanced nanopore system for investigating various functional nanostructures, we analysed the translocation of proteins and protein spherical nucleic acids (ProSNAs). ProSNAs are a new class of “soft” nanoparticles with wide-spread applications in bottom-up materials assembly, intracellular sensing, and CRISPR-based gene editing (38-40). ProSNAs are based on the spherical nucleic acid (SNA) architecture and consist of a protein core functionalized with a dense shell of DNA strands (38). Compared to its analogous native protein, ProSNAs, i.e. the protein functionalized with ssDNA (single-stranded DNA), show an increase in cellular uptake, with the oligonucleotide density playing a critical role in the cellular uptake efficiency (40). Hence, a rational investigation of the DNA-mediated functionalization of these hybrid nanostructures can support the development of new architectures. Here, we used β-galactosidase (β-gal) ProSNAs thanks to their well-established stability and ease of synthesis (Section S13). As depicted in Figure 5A, our polymer electrolyte enhanced sensing approach enabled the detection of the native β-gal protein and β-gal SNAs with two different DNA loadings: 22 ssDNAs (β-gal SNA_22_) and 42 ssDNAs (β-gal SNA_42_), respectively. The average conductive peak current obtained for the native β-gal protein was centred at 32 pA, whilst a substantial increase was observed for the β-gal SNA samples with a peak centred at 94 pA for the β-gal SNA_22_ and 171 pA for β -gal SNA_42_ (Figure 5B-C). Similarly, an increase in the translocation duration was observed between the native β-gal protein and β-gal SNAs (Table S3 and Figure S17). Importantly, apart from the clear discrimination between the native protein and the DNA-functionalized analogous, we were also able to differentiate the β-gal SNAs with two different DNA loadings (SNA_22_ and SNA_42_). The increase in the density of the functionalized ssDNA can change the effective size of the particle and lead to an increase in the hydrodynamic radius. Remarkably, even at low DNA loading (SNA_22_), comprising only several sparsely attached ssDNA to the protein surface, we were able to provide a clear discrimination from the bare protein serving as scaffold for attachment. In similar fashion, we showed the ability to differentiate between the two different ssDNA loadings, with a proportional increase in the peak current and dwell time with the increasing density of the ssDNAs attached to the protein. These results are in line with the approximately two-fold increase in the DNA shell density of the functionalized protein. This rapid yet easy-to-implement nanopore sensing allowed us to probe small variations in the density of the soft DNA shell on the surface of the protein. Taken together, these results show that the polymer-electrolyte nanopore system can enable the characterization of soft nanomaterials and allow for the discrimination of nanoparticles based on their functionalization state. These measurements provide a basis to investigate functionalization strategies with biomolecules (i.e., different densities/moieties) for hybrid materials at the nanoscale level with implications in cellular imaging, sensing, or drug delivery (41,42).

**FIGURE 5.**
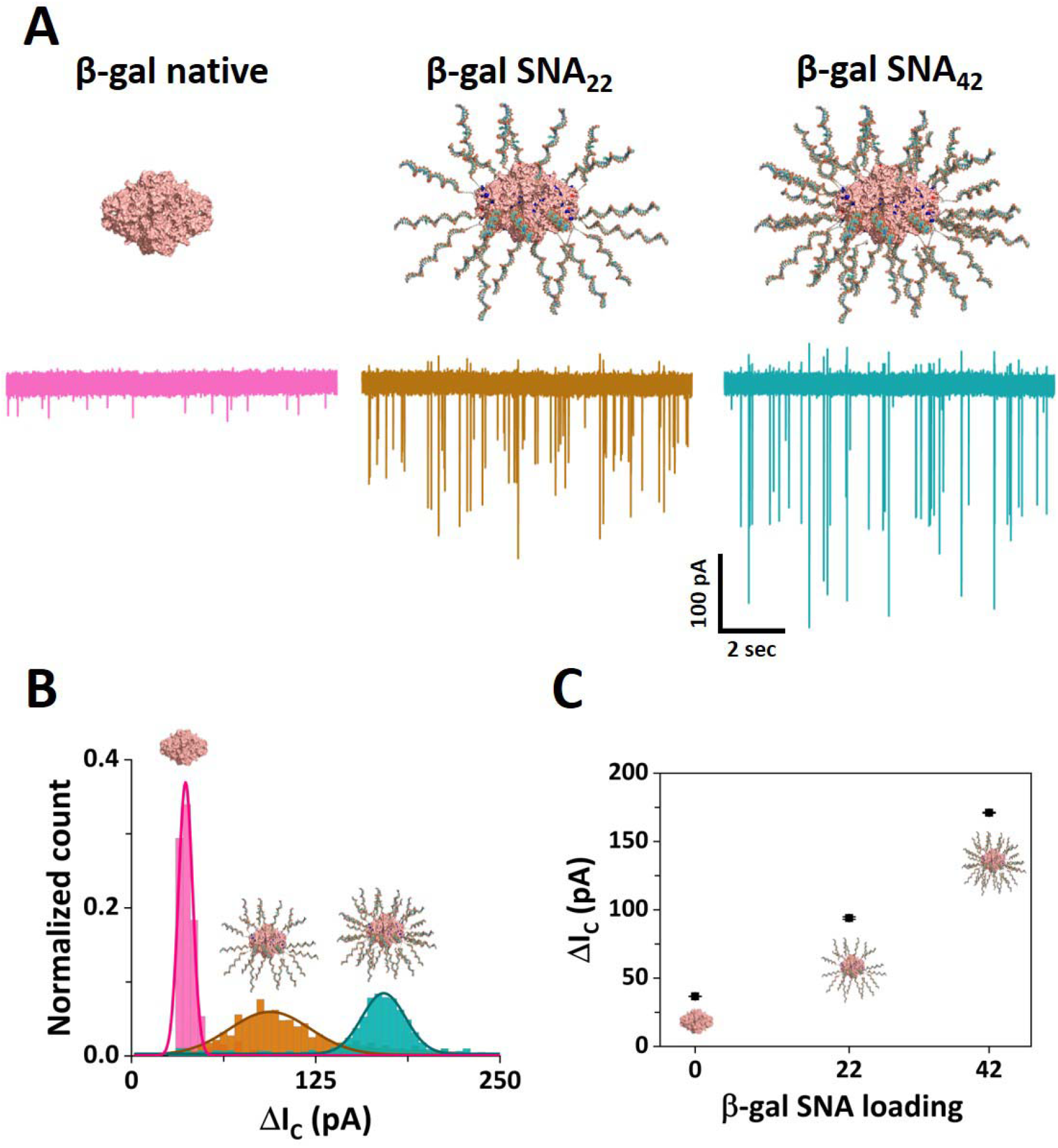
Nanopore sensing of protein spherical nucleic acids **(A)** Nanopore translocation ion current traces obtained for the native β-gal (left), β-gal SNA_22_ (middle), and β-gal SNA_42_ (right). The current and time scales are the same of all the ion current traces. **(B)** Histograms of the conductive peak current for native β-gal (pink bars), β-gal SNA_22_ (brown bars), and β-gal SNA_42_ (teal bars). **(C)** Average conductive peak current as a function of the oligonucleotide loading. The error bars represent the standard error. Nanopore measurements were carried in 50% PEG and 100 mM KCl using a 30 nm diameter pore size biased at −700 mV.

## CONCLUSIONS

We have demonstrated the development and implementation of a nanopore system enhanced by a polymer electrolyte enabling the analytical characterization of nanoparticles. Using experiments and finite element modelling, we provided a mechanistic description of the ion current signals and further employed the system to analyse heterogenous gold nanoparticle mixtures. We then demonstrated the unique ability of our approach to fingerprint nanoparticle samples at low ionic strength (10 mM) and exemplify the power of the system by characterizing the degree of branching of “hard” anisotropic Au nanostars and the nucleic acid coverage of “soft” ProSNAs. This universal system will complement the toolbox of nanomaterials characterization technique and enable the real-time optimization of flow synthesis of a wide range of nanomaterials.

## EXPERIMENTAL SECTION

### Chemicals and materials

All reagents used in the translocation experiments were prepared using ultra-pure water (18.2 MΩ.cm) from Millipore system and further filtered through a 0.22 μm syringe. KCl, Triton-X, EDTA, and PEG reagents were purchased from Sigma Aldrich. Gold PEG carboxyl-capped nanospheres (10, 20, 30, 40, and 50 nm diameter), silver citrate-capped nanospheres (30 nm diameter), and platinum citrate-capped nanospheres (30 nm nanospheres) were purchased from NanoComposix. Silver wire (0.25 mm diameter) used in the nanopore experiments were obtained from Alfa Aesar.

### Standard nanoparticles characterization

The stability of the gold nanoparticles diluted in the KCl translocation buffer was probed by UV-Vis measurements (Figure S2) using a NanoDrop ND-1000 spectrophotometer (Thermo Scientific). The size distribution of the standard nanoparticles in solution was determined by Zetasizer NanoZS (Malvern Instruments Ltd.) (Figure S2). All the standard nanosphere samples were used as received.

### Nanopore fabrication and characterization

The nanopores were fabricated starting from 1.0 mm x 0.5 mm quartz capillaries (QF120-90-10; Sutter Instrument, UK) with the SU-P2000 laser puller (World Precision Instruments, UK), using a two-line program: (1) HEAT, 750; FILAMENT, 4, VELOCITY, 30; DELAY, 145, PULL, 80; (2) HEAT, 600, FILAMENT, 3; VELOCITY, 40; DELAY, 135; PULL, 150. The pulling parameters are instrument specific and lead to glass nanopore with a diameter of approximately 60 nm. Adjustments of the HEAT AND PULL parameters were made to fabricate other pore sizes specified in this study. The pulled glass nanopores were characterized by measuring their pore resistance in 0.1 M KCl and the pore dimensions were confirmed by Scanning Electron Microscopy (SEM) using a Nova NanoSEM at an accelerating voltage of 3–5 kV. The characterization of 60 nm glass nanopore is exemplified in Figure S1.

### Nanopore translocation measurements

Unless otherwise specified, the translocation experiments were carried out by filling the glass nanopore with the translocation buffer (50 mM KCl, 0.01% Triton-X, 10 mM Tris, 1 mM EDTA, pH 8.0) containing the nanoparticles. The glass nanopore was then immersed in a similar buffer with the addition of 50% (w/v) Polyethylene Glycol (PEG) 35 kDa (ultrapure grade, Sigma Aldrich). The notation of the 50% PEG in the text refers to the utilization of 50% (w/v) Polyethylene Glycol (PEG) 35 kDa. An Ag/AgCl wire (0.25 mm diameter, GoodFellow UK) was inserted in the glass nanopore barrel and acted as the working electrode, while a second Ag/AgCl wire was immersed in the bath and acted as the counter and reference electrodes. The nanoparticles were driven from inside the glass nanopore into the external bath by applying a negative potential to the working electrode placed inside the glass nanopore with respect to the reference electrode in the bath. The ion current was recorded with a MultiClamp 700B patch-clamp amplifier (Molecular Devices) in voltage-clamp mode. Data was acquired at a 100 kHz sampling rate with a 20 kHz low-pass filter using the pClamp10 software (Molecular Devices). The ion current traces were further analyzed with the MATLAB script Transalyzer, developed by Plesa et al. (43). The obtained translocation events were analyzed by applying a 7-sigma threshold level from the baseline, and only the events above the threshold were considered as translocation events (Figure S4). The obtained events were further analyzed and plotted using Origin 2019b.

### Numerical simulations

To provide a mechanistic understanding of the experimentally observed current responses during nanoparticle translocation in our system, we developed numerical simulations describing the electric potential, ion concentrations, and fluid flow within and around the aperture of the glass nanopore. The simulations, which we describe briefly below, are based on a model we developed previously to describe a glass nanopore immersed in a polymer electrolyte (32). For a more detailed description, including a full list of parameters and boundary conditions, see Supporting Information, section S16 (32). The commercial finite element software COMSOL Multiphysics (version 5.6) was used to solve the equations describing the spatially varying quantities listed above. Boundary conditions were selected to reflect the experimental system, including the surface charge on the glass/nanoparticle and the bulk solution concentrations. K^+^ and Cl^-^ transport depends on the phase (PEG+KCl or KCl) with diffusion coefficients chosen to match the experimentally measured solution conductivities. In KCl, the diffusion coefficients of the K^+^ and Cl^-^ are approximately equal (44) (D_K+_:D_Cl-_ = 0.49:0.51) while in the PEG D_K+_:D_Cl-_ = 0.35:0.65 ratio, as previously determined (32). The latter reflects cation affinity to PEG. For simplicity, the interface between the PEG+KCl and KCl was taken to have zero width, i.e., mixing of the solutions was neglected. In the absence of a nanoparticle, the interface was taken to be the disk exactly at the mouth of the pipette, while a nanoparticle translocation was taken to perturb this interface outward (see Figure S18). The current was calculated from integrating the ion flux at the top of the nanopore.

### Gold nanostars synthesis

First, citrate-capped gold nanoparticle (12 nm) seeds were synthesized using the Turkevich method (45).⍰Briefly, 100 mL ultra-pure boiling water and 200 μL of 0.5 M HAuCl_4_⍰were dispersed for 10 s. 15 mL of 1% trisodium citrate was added. After the final colour change, the solution was boiled for 15-30 minutes. Then the solution was cooled, filtered through a 0.22 μm nitrocellulose membrane, and stored at 4⍰°C until further use. The Au nanostars synthesis followed the Vo-Dinh group surfactant-free procedure with a few modifications (34).⍰In brief, under room temperature and fast stirring, 10 mL of ultra-pure H_2_O, 10 μL 1 M HCl, and 493 μL of 0.5 M HAuCl_4_⍰were added to a plastic scintillation vial. Then, 100 μL of Au seeds were added, and 10 seconds afterward quickly added AgNO_3_⍰of various concentrations (1-3 mM; samples named S5, S10, and S30 relating to the AgNO_3_⍰final concentration, 5 μM, 10 μM, and 30 μM, respectively) and 50 μL of 0.1 M AA, simultaneously. Finally, 10 s afterward, transferred the stirred NS solution and immediately used. After the Au nanostar synthesis, 325 μL of thiol stabilizer (HS-PEG-COOH) was added to the Au nanostars, vortexed, then idle for 30 min at room temperature. After 30 minutes, the solution was centrifuged and washed 3 times with ultra-pure H_2_O (1500 g, 15 min, 4°C). Lastly, the samples were redispersed in ultra-pure H_2_O and stored at 4 °C until used.

### Gold nanostars characterization

Au Nanostars with thiol stabilizer were prepared for imaging by dropping 10 μL of 2-fold diluted sample onto a 200-mesh grid. The sample was allowed to dry overnight at room temperature. Then transmission electron microscope (TEM; Hitachi H7650) imaging was performed at 80 kV with an AMT BIOSPR16 camera (Figure S13). To determine the size and concentration, the samples were measured via NanoSight NS300 nanoparticle tracking analysis (NTA) system with a 532 nm laser. 20-, 40-, and 80-fold dilutions of the samples were used for the measurements. Extinction spectra of the samples were collected via a Biotek Microplate reader (Figure 12).

### DNA synthesis and characterization

All oligonucleotides used in this work (Table S2) were synthesized using a MerMade 6 instrument (LGC, Biosearch Technologies) at 1 μmol scale. Reagents for DNA synthesis were purchased from Glen Research. Controlled pore glass (CPG) beads (Glen Research, Cat. No. 20-5041-10) were used as the solid support for DNA synthesis. The synthesized products were cleaved from the CPG beads and deprotected using 0.5 mL of 30% ammonium hydroxide (Thermo Fisher, Cat. No. A669) for 16 h at room temperature. After 16 h, the solution was run through a NAP-5 column (Cytiva illustra, Cat. No. 17085301) following manufacturer’s protocol. The eluate was purified using reversed phase HPLC (Vanquish HPLC, Thermo Fisher, USA) using a C18 column (Thermo Fisher, Cat. No. 41005-259070A). A gradient of 0 to 70% A→B over 40 minutes was used. A was 30 mM triethylammonium acetate (Thermo Fisher, Cat. No. O4885-1) with 3% (v/v) acetonitrile (Thermo Fisher, Cat. No. A996SK-4) and B was 100% acetonitrile. The collected fractions were lyophilized and redissolved in water. The mass of the oligonucleotides was determined using matrix-assisted laser desorption ionization time-of-flight mass spectrometry (MALDI-TOF MS, Bruker, AutoFlex Max) using 2’,6’-dihydroxyacetophenone (DHAP, Sigma, Cat. No. 37468) as a matrix. The absorbance (A*λ*) of oligonucleotides was measured as a function of wavelength (*λ*) using UV-Vis spectroscopy (Agilent, Cary 60 UV-Vis Spectrophotometer) using a quartz cuvette (Thermo Fisher, Cat. No. 50-753-2877) with 1 cm pathlength. Using the extinction coefficient (*ελ*) at 260 nm (obtained from the IDT Oligo Analyzer tool), the concentration (*c*) of the purified products was determined using Beer’s law (Equation 1):

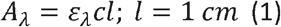

### ProSNA synthesis and characterization

β-gal SNAs were synthesized by following previously established procedures with some modifications (S13) (38,39). Briefly, surface cysteine groups are first modified with Alexa Fluor 647 C_2_ maleimide. Following this step, NHS-PEG_4_-azides are conjugated to the surface-accessible lysine residues. Finally, DBCO-terminated DNA is attached to the azide-modified proteins through copper-free click chemistry. Details of the synthesis steps are described in Supporting Information, section S13. The synthesis of β-gal SNAs was confirmed through UV-Vis and native PAGE gel electrophoresis, as shown in Figure S15 and Figure S16.

## DECLARATION OF INTERESTS

The authors declare no competing interests.

## ACKNOWLEDGEMENTS

The authors gratefully acknowledge Dr. Zabeada Aslam and the Leeds Electron Microscopy and Spectroscopy centre (LEMAS) for their support and assistance with TEM characterization of the gold nanoparticles. We thank Dr. Alexander Kulak for the help provided with SEM imaging of the glass nanopores. S.C. and P.A. acknowledge funding from the European Union’s Horizon 2020 research and innovation 7 program under the Marie Skłodowska-Curie MSCA-ITN grant agreement no. <u>812398</u>, through the single-entity nanoelectrochemistry, SENTINEL, project. P.A. Acknowledge funding from the European Research Council (ERC) under the project DYNAMIN, grant agreement no. <u>788968</u>. P.A and C.W. acknowledge funding from the Engineering and Physical Sciences Research Council UK (EPSRC) Healthcare Technologies for the grant <u>EP/W004933/1</u>, and S.C. and C.W. acknowledges funding from the Medical Research Council (MRC) UK under the grant number <u>MR/N029976/1</u>. T.I.P. acknowledges the University of Leeds Faculty of Biological Sciences and the US Dept of Education (USDE) for funding. Schematics used in our figures were adapted from BioRender.com.

## DATA AVAILABILITY

## Notes

### Competing Interest Statement

The authors have declared no competing interest.

### Summary of Updates

Updated the name of one of the authors that has been misspelled.

## REFERENCES

1. Zhang, H. 2020. Molecularly Imprinted Nanoparticles for Biomedical Applications. Advanced Materials. 32(3):1806328, doi: https://doi.org/10.1002/adma.201806328.

2. Tan, H. W., J. An, C. K. Chua, and T. Tran. 2019. Metallic Nanoparticle Inks for 3D Printing of Electronics. Advanced Electronic Materials. 5(5):1800831, doi: https://doi.org/10.1002/aelm.201800831.

3. Wang, L., M. Hasanzadeh Kafshgari, and M. Meunier. 2020. Optical Properties and Applications of Plasmonic-Metal Nanoparticles. Advanced Functional Materials. 30(51):2005400, doi: https://doi.org/10.1002/adfm.202005400.

4. Stark, W. J., P. R. Stoessel, W. Wohlleben, and A. Hafner. 2015. Industrial applications of nanoparticles. Chemical Society Reviews. 44(16):5793–5805, doi: 10.1039/C4CS00362D.

5. Xu, X., D. Valavanis, P. Ciocci, S. Confederat, F. Marcuccio, J.-F. Lemineur, P. Actis, F. Kanoufi, and P. R. Unwin. 2023. The New Era of High-Throughput Nanoelectrochemistry. Analytical Chemistry. 95(1):319–356, doi: 10.1021/acs.analchem.2c05105.

6. Tomaszewska, E., K. Soliwoda, K. Kadziola, B. Tkacz-Szczesna, G. Celichowski, M. Cichomski, W. Szmaja, and J. Grobelny. 2013. Detection limits of DLS and UV-Vis spectroscopy in characterization of polydisperse nanoparticles colloids. Journal of Nanomaterials. 2013:60–60

7. Filipe, V., A. Hawe, and W. Jiskoot. 2010. Critical evaluation of Nanoparticle Tracking Analysis (NTA) by NanoSight for the measurement of nanoparticles and protein aggregates. Pharmaceutical research. 27:796–810

8. Modena, M. M., B. Rühle, T. P. Burg, and S. Wuttke. 2019. Nanoparticle Characterization: What to Measure? Advanced Materials. 31(32):1901556, doi: https://doi.org/10.1002/adma.201901556.

9. Miles, B. N., A. P. Ivanov, K. A. Wilson, F. Doğan, D. Japrung, and J. B. Edel. 2013. Single molecule sensing with solid-state nanopores: novel materials, methods, and applications. Chemical Society Reviews. 42(1):15–28

10. Lu, S.-M., Y.-Y. Peng, Y.-L. Ying, and Y.-T. Long. 2020. Electrochemical sensing at a confined space. Analytical chemistry. 92(8):5621–5644

11. Waduge, P., R. Hu, P. Bandarkar, H. Yamazaki, B. Cressiot, Q. Zhao, P. C. Whitford, and M. Wanunu. 2017. Nanopore-based measurements of protein size, fluctuations, and conformational changes. ACS nano. 11(6):5706–5716

12. Houghtaling, J., C. Ying, O. M. Eggenberger, A. Fennouri, S. Nandivada, M. Acharjee, J. Li, A. R. Hall, and M. Mayer. 2019. Estimation of Shape, Volume, and Dipole Moment of Individual Proteins Freely Transiting a Synthetic Nanopore. ACS Nano. 13(5):5231–5242, doi: 10.1021/acsnano.8b09555.

13. Arjmandi, N., W. Van Roy, L. Lagae, and G. Borghs. 2012. Measuring the Electric Charge and Zeta Potential of Nanometer-Sized Objects Using Pyramidal-Shaped Nanopores. Analytical Chemistry. 84(20):8490–8496, doi: 10.1021/ac300705z.

14. Lee, K., K.-B. Park, H.-J. Kim, J.-S. Yu, H. Chae, H.-M. Kim, and K.-B. Kim. 2018. Recent Progress in Solid-State Nanopores. Advanced Materials. 30(42):1704680, doi: https://doi.org/10.1002/adma.201704680.

15. Yang, W., B. Radha, A. Choudhary, Y. You, G. Mettela, A. K. Geim, A. Aksimentiev, A. Keerthi, and C. Dekker. 2021. Translocation of DNA through Ultrathin Nanoslits. Advanced Materials. 33(11):2007682, doi: https://doi.org/10.1002/adma.202007682.

16. Ren, R., M. Sun, P. Goel, S. Cai, N. A. Kotov, H. Kuang, C. Xu, A. P. Ivanov, and J. B. Edel. 2021. Single-Molecule Binding Assay Using Nanopores and Dimeric NP Conjugates. Advanced Materials. 33(38):2103067, doi: https://doi.org/10.1002/adma.202103067.

17. German, S. R., L. Luo, H. S. White, and T. L. Mega. 2013. Controlling Nanoparticle Dynamics in Conical Nanopores. The Journal of Physical Chemistry C. 117(1):703–711, doi: 10.1021/jp310513v.

18. Smeets, R. M. M., U. F. Keyser, D. Krapf, M.-Y. Wu, N. H. Dekker, and C. Dekker. 2006. Salt Dependence of Ion Transport and DNA Translocation through Solid-State Nanopores. Nano Letters. 6(1):89–95, doi: 10.1021/nl052107w.

19. Kowalczyk, S. W., D. B. Wells, A. Aksimentiev, and C. Dekker. 2012. Slowing down DNA Translocation through a Nanopore in Lithium Chloride. Nano Letters. 12(2):1038–1044, doi: 10.1021/nl204273h.

20. Platt, M., G. R. Willmott, and G. U. Lee. 2012. Resistive Pulse Sensing of Analyte-Induced Multicomponent Rod Aggregation Using Tunable Pores. Small. 8(15):2436–2444, doi: https://doi.org/10.1002/smll.201200058.

21. Maugi, R., P. Hauer, J. Bowen, E. Ashman, E. Hunsicker, and M. Platt. 2020. A methodology for characterising nanoparticle size and shape using nanopores. Nanoscale. 12(1):262–270, doi: 10.1039/C9NR09100A.

22. Chau, C., F. Marcuccio, D. Soulias, M. A. Edwards, A. Tuplin, S. E. Radford, E. Hewitt, and P. Actis. 2022. Probing RNA Conformations Using a Polymer–Electrolyte Solid-State Nanopore. ACS Nano. 16(12):20075–20085, doi: 10.1021/acsnano.2c08312.

23. Chau, C. C., S. E. Radford, E. W. Hewitt, and P. Actis. 2020. Macromolecular Crowding Enhances the Detection of DNA and Proteins by a Solid-State Nanopore. Nano Letters. 20(7):5553–5561, doi: 10.1021/acs.nanolett.0c02246.

24. Prabhu, A. S., T. Z. N. Jubery, K. J. Freedman, R. Mulero, P. Dutta, and M. J. Kim. 2010. Chemically modified solid state nanopores for high throughput nanoparticle separation. Journal of Physics: Condensed Matter. 22(45):454107, doi: 10.1088/0953-8984/22/45/454107.

25. Wen, C., S. Zeng, Z. Zhang, and S.-L. Zhang. 2018. Group Behavior of Nanoparticles Translocating Multiple Nanopores. Analytical Chemistry. 90(22):13483–13490, doi: 10.1021/acs.analchem.8b03408.

26. Lan, W.-J., D. A. Holden, B. Zhang, and H. S. White. 2011. Nanoparticle Transport in Conical-Shaped Nanopores. Analytical Chemistry. 83(10):3840–3847, doi: 10.1021/ac200312n.

27. Terejánszky, P., I. Makra, P. Fürjes, and R. E. Gyurcsányi. 2014. Calibration-Less Sizing and Quantitation of Polymeric Nanoparticles and Viruses with Quartz Nanopipets. Analytical Chemistry. 86(10):4688–4697, doi: 10.1021/ac500184z.

28. Sensale, S., Z. Peng, and H.-C. Chang. 2019. Biphasic signals during nanopore translocation of DNA and nanoparticles due to strong ion cloud deformation. Nanoscale. 11(47):22772–22779, doi: 10.1039/C9NR05223B.

29. Chen, K., L. Shan, S. He, G. Hu, Y. Meng, and Y. Tian. 2015. Biphasic Resistive Pulses and Ion Concentration Modulation during Particle Translocation through Cylindrical Nanopores. The Journal of Physical Chemistry C. 119(15):8329–8335, doi: 10.1021/acs.jpcc.5b00047.

30. Menestrina, J., C. Yang, M. Schiel, I. Vlassiouk, and Z. S. Siwy. 2014. Charged Particles Modulate Local Ionic Concentrations and Cause Formation of Positive Peaks in Resistive-Pulse-Based Detection. The Journal of Physical Chemistry C. 118(5):2391–2398, doi: 10.1021/jp412135v.

31. Lastra, L. S., Y. M. N. D. Y. Bandara, M. Nguyen, N. Farajpour, and K. J. Freedman. 2022. On the origins of conductive pulse sensing inside a nanopore. Nature Communications. 13(1):2186, doi: 10.1038/s41467-022-29758-8.

32. Marcuccio, F., D. Soulias, C. C. C. Chau, S. E. Radford, E. Hewitt, P. Actis, and M. A. Edwards. 2023. Mechanistic Study of the Conductance and Enhanced Single-Molecule Detection in a Polymer–Electrolyte Nanopore. ACS Nanoscience Au. doi: 10.1021/acsnanoscienceau.2c00050.

33. Wei, X., A. Popov, and R. Hernandez. 2022. Electric Potential of Citrate-Capped Gold Nanoparticles Is Affected by Poly(allylamine hydrochloride) and Salt Concentration. ACS Applied Materials & Interfaces. 14(10):12538–12550, doi: 10.1021/acsami.1c24526.

34. Yuan, H., C. G. Khoury, H. Hwang, C. M. Wilson, G. A. Grant, and T. Vo-Dinh. 2012. Gold nanostars: surfactant-free synthesis, 3D modelling, and two-photon photoluminescence imaging. Nanotechnology. 23(7):075102

35. Strobbia, P., V. Cupil-Garcia, B. M. Crawford, A. M. Fales, T. J. Pfefer, Y. Liu, M. Maiwald, B. Sumpf, and T. Vo-Dinh. 2021. Accurate in vivo tumor detection using plasmonic-enhanced shifted-excitation Raman difference spectroscopy (SERDS). Theranostics. 11(9):4090

36. Indrasekara, A. S. D. S., S. Meyers, S. Shubeita, L. C. Feldman, T. Gustafsson, and L. Fabris. 2014. Gold nanostar substrates for SERS-based chemical sensing in the femtomolar regime. Nanoscale. 6(15):8891–8899

37. Barbosa, S., A. Agrawal, L. Rodríguez-Lorenzo, I. Pastoriza-Santos, R. A. Alvarez-Puebla, A. Kornowski, H. Weller, and L. M. Liz-Marzán. 2010. Tuning Size and Sensing Properties in Colloidal Gold Nanostars. Langmuir. 26(18):14943–14950, doi: 10.1021/la102559e.

38. Ebrahimi, S. B., D. Samanta, C. D. Kusmierz, and C. A. Mirkin. 2022. Protein transfection via spherical nucleic acids. Nature Protocols. 17(2):327–357, doi: 10.1038/s41596-021-00642-x.

39. Samanta, D., S. B. Ebrahimi, C. D. Kusmierz, H. F. Cheng, and C. A. Mirkin. 2020. Protein Spherical Nucleic Acids for Live-Cell Chemical Analysis. Journal of the American Chemical Society. 142(31):13350–13355, doi: 10.1021/jacs.0c06866.

40. Samanta, D., S. B. Ebrahimi, and C. A. Mirkin. 2020. Nucleic-Acid Structures as Intracellular Probes for Live Cells. Advanced Materials. 32(13):1901743, doi: https://doi.org/10.1002/adma.201901743.

41. Huang, X., et al. 2021. DNA scaffolds enable efficient and tunable functionalization of biomaterials for immune cell modulation. Nature Nanotechnology. 16(2):214–223, doi: 10.1038/s41565-020-00813-z.

42. He, L., J. Mu, O. Gang, and X. Chen. 2021. Rationally programming nanomaterials with DNA for biomedical applications. Advanced Science. 8(8):2003775

43. Plesa, C., and C. Dekker. 2015. Data analysis methods for solid-state nanopores. Nanotechnology. 26(8):084003

44. Haynes, W. M. 2016. CRC handbook of chemistry and physics 97th Edition. CRC press.

45. Turkevich, J., P. C. Stevenson, and J. Hillier. 1951. A study of the nucleation and growth processes in the synthesis of colloidal gold. Discussions of the Faraday Society. 11:55–75

