## Supporting Information for "Next-Generation Nanopore Sensors for Enhanced Detection of Nanoparticles"

### **S1: Nanopore characterization.**

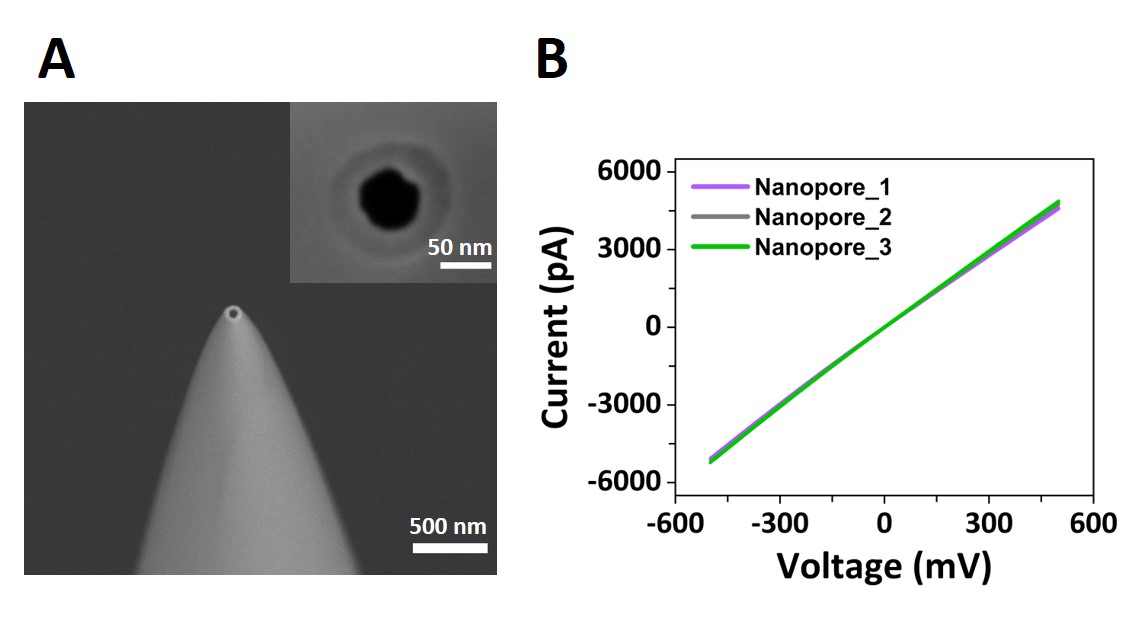

**Figure S1: (A)** Scanning electron micrograph of a representative glass nanopores used in this study. Inset: top view of the nanopore tip showing that it has a ~60 nm internal diameter. Note: The micrograph foreshortens depth resulting in an exaggerated angle of the pipette wall **(B)** Current-voltage curves (I-V) of three glass nanopores recorded in 0.1 M KCl. All three pipettes were pulled with an identical program which gave ~60 nm pores, such as the one shown in panel A.

### **S2: AuNP stability assessment in 100 mM KCl.**

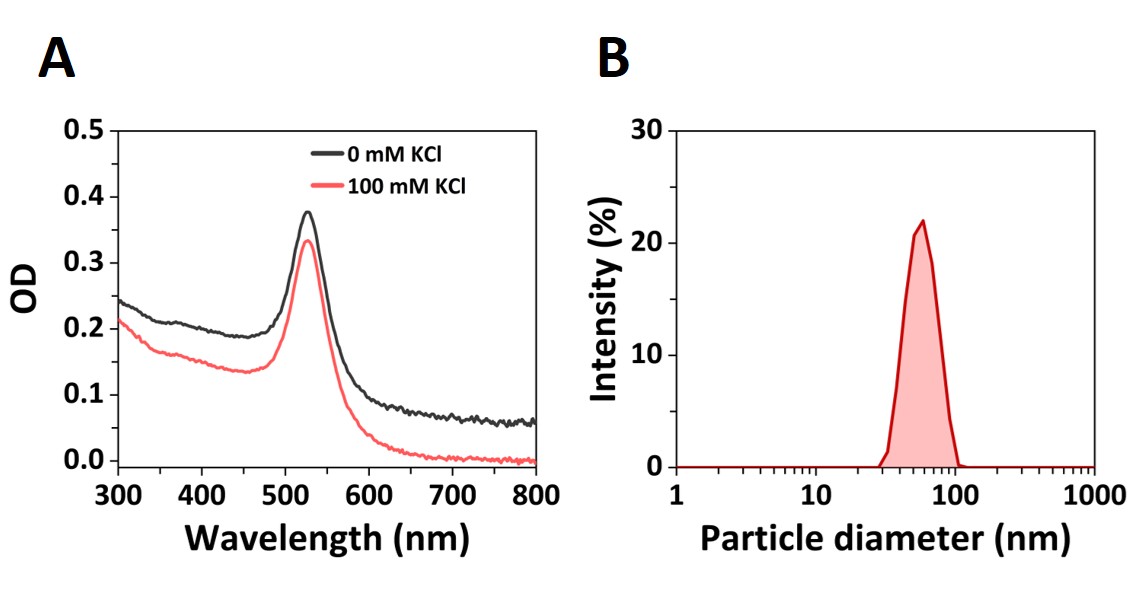

**Figure S2:** Stability assessment AuNP suspended in 100 mM KCl buffer solution. **(A)** UV-Vis spectra of 50 nm PEG carboxyl-capped AuNPs (25 µg/mL) in 100 mM KCl buffer solution measured after 1 hr incubation (red line) compared with a preparation in Milli-Q water (black line). A similar spectrum is observed in both cases indicating that the nanoparticles in 100 mM KCl buffer solution are stable for >1 hr. All measurements of samples in this work were completed within 1 hr, thus there is no concern over the stability of the particles. The peak in optical absorbance at 520 nm is due to a plasmon resonance. No broadening or shift to higher wavelengths of the absorbance peak was observed in 100 mM KCl, suggesting minimal aggregation occurring. **(B)** Dynamic light scattering (DLS) measured size distribution (hydrodynamic diameter) of nominally 50 nm diameter PEG carboxyl-capped AuNPs (25 µg/mL) measured in 100 mM KCl after 1hr incubation. The size distribution measured shows the presence of a homogenous suspension of nanoparticles without aggregates formation.

### **S3: Enhancement of AuNP resistive pulses using the polymer electrolyte.**

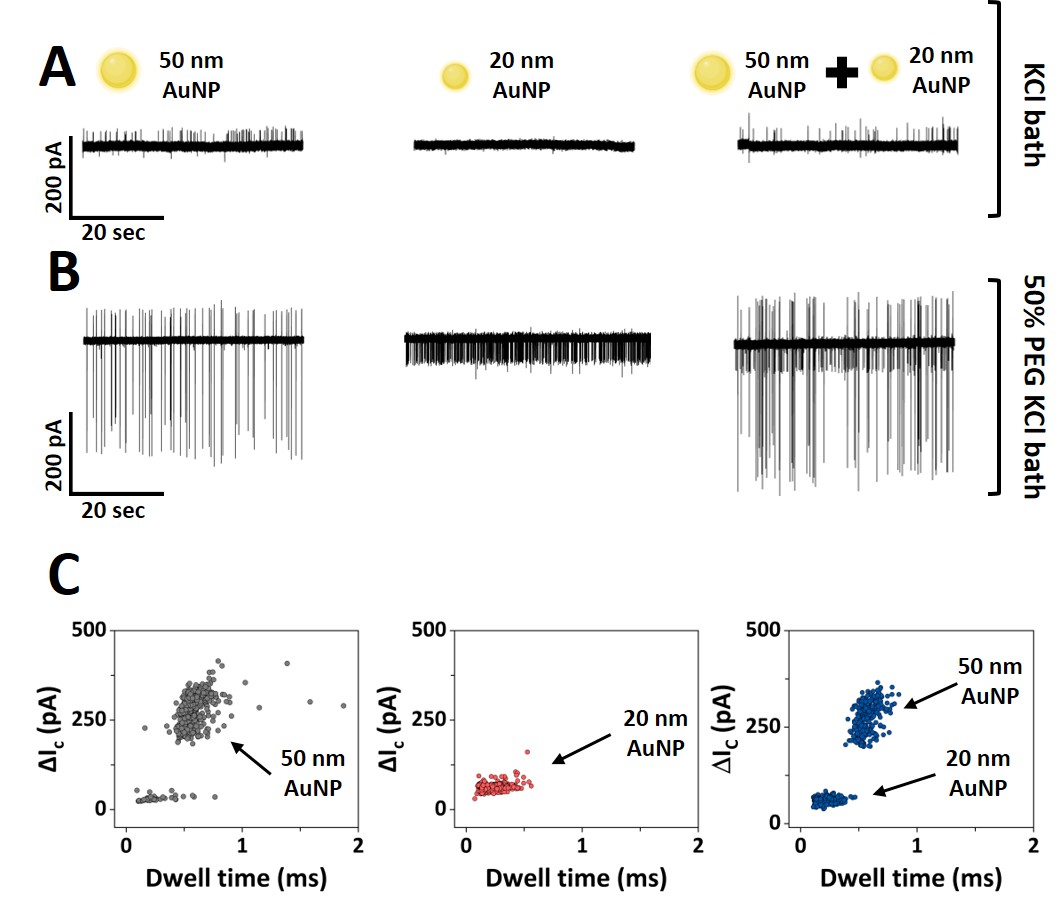

**Figure S3:** Ion current traces of nanoparticle translocations through a ~60 nm diameter pore in which the external electrolyte bath was either **(A)** 50 mM KCl or **(B)** 50 mM KCl + 50% PEG. The inside of the nanopore contained PEG carboxyl-capped AuNPs of 50 nm diameter (left), 20 nm diameter (middle), or a mixture of a 50 nm and 20 nm diameter (right). A single glass nanopores (~60 nm diameter) was used for each nanoparticle mixture in the two conditions. In the absence of PEG in the bath solution 20 nm AuNPs translocation peaks are not resolved from the current baseline. The current and time scales are the same for all the ion current traces in panels A and B. In all cases, the glass nanopore contained 50 mM KCl without PEG. An approximately two-fold decrease in the event frequency was observed between the KCl and KCl + 50% PEG external bath for the 50 nm AuNPs translocation, attributed to the presence of the viscous external bath when 50% PEG is added to the electrolyte bath. **(C)** Scatter plots of the conductive peak current (ΔI_C_) versus dwell time of the translocation events extracted from the ion current traces in panel (B). Ion current traces were recorded at -500 mV. The concentration of the individual AuNPs (50 nm and 20 nm AuNPs) was 25 µg/mL, whereas the 50 nm and 20 nm AuNP mixture was recorded at mixture ratio of 10:1 50-to-20 nm AuNPs (particle concentration in the mixture: 1.89 x 10^10^ particles/mL for 50 nm AuNPs and 3.1x 10^9^ particles/mL for 20 nm AuNPs) to account for the difference in translocation frequency of 20nm and 50 nm AuNPs at the same mass concentration (µg/mL). Optimization of the event rate has not been a focus in this study.

### **S4: Detection of translocation events.**

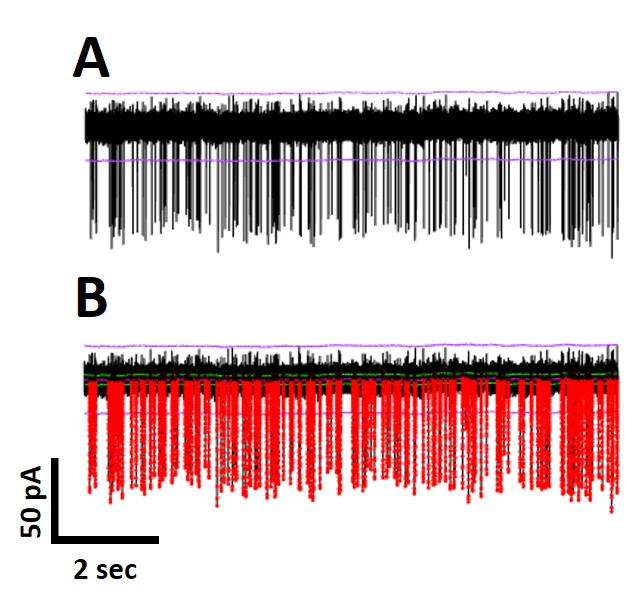

**Figure S4: (A)** Representative ion current trace (20 nm diameter PEG carboxyl-capped AuNPs at concentration of 25 µg/mL, -500 mV, with 50% PEG in 50 mM KCl external bath). Purple lines indicate the ±7σ threshold used to detect AuNP translocations. **(B)** Same trace with the perturbations in the ion current >7σ highlighted in red. AuNPs translocation events were assigned using the Transalyzer Matlab script (1).

### **S5: Control experiments for nanopore translocation.**

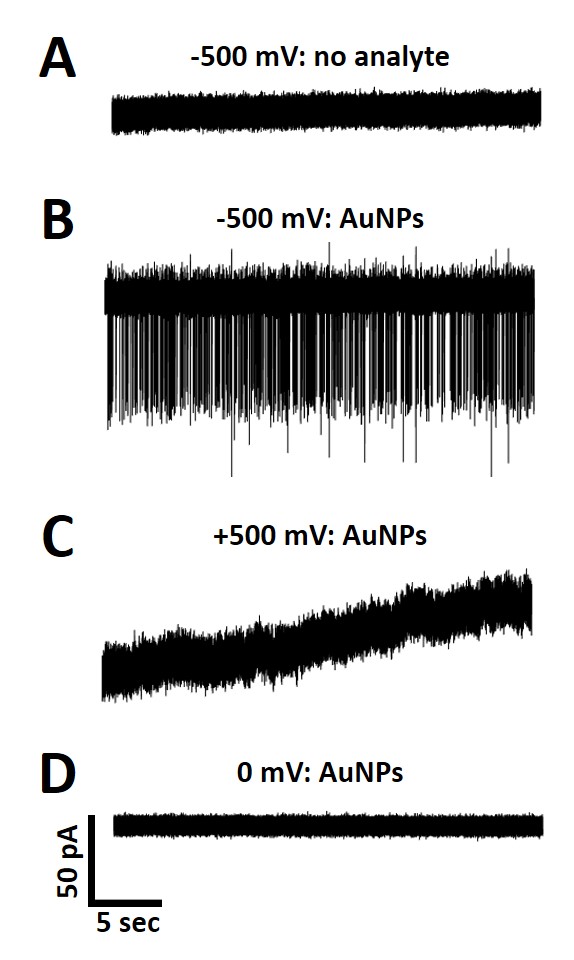

**Figure S5:** Ion current traces recorded with a 60 nm diameter glass nanopore in a bath solution containing 50% PEG in 50 mM KCl. The glass nanopore was biased at **(A)** -500 mV when no analyte was added inside the glass nanopore; **(B)** -500 mV when 20 nm diameter PEG carboxyl-capped AuNPs were added inside the glass nanopore; **(C)** +500 mV when 20 nm diameter AuNPs were added inside the glass nanopore; **(D)** 0 mV when 20 nm diameter AuNPs were added inside the glass nanopore. The current and timescales are the same for all graphs. A concentration of 25 µg/mL was used for the recordings with 20 nm diameter AuNPs.

### **S6: Characteristics of AuNP translocations.**

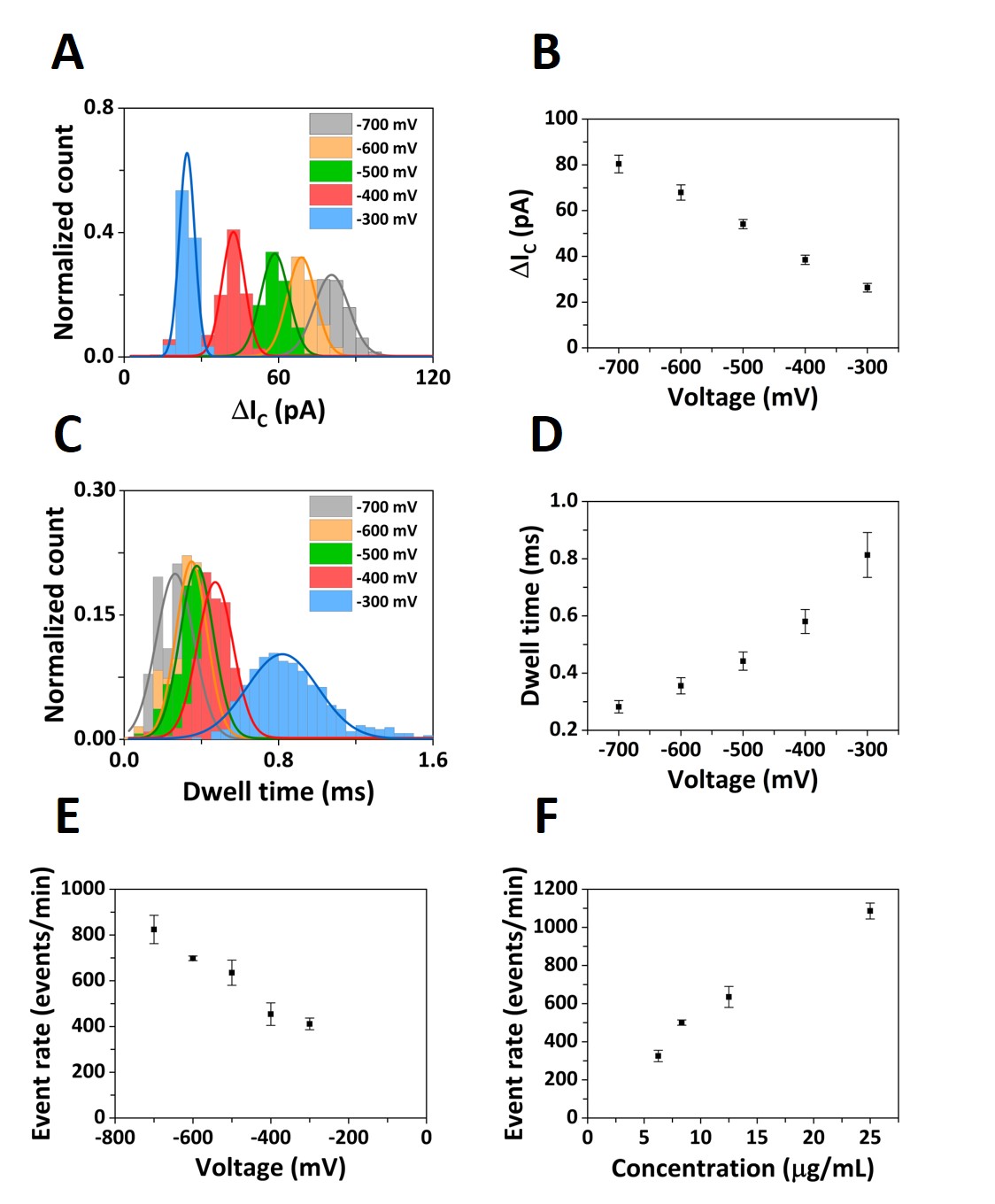

**Figure S6:** Translocation event characteristics of 20 nm diameter PEG carboxyl-capped AuNPs translocating through a ~60 nm diameter pore with 50% PEG in 50 mM KCl external bath. Histograms of the conductive peak current **(A)** and dwell time **(C)** distributions for applied voltages *V* = 300 mV, ‑400 mV, -500 mV, -600 mV, and -700 mV. The solid lines represent Gaussian fits to the distributions. Average conductive peak current **(B)** and dwell time **(D)** at *V* = -300 mV, -400 mV, -500 mV, -600 mV, and -700 mV. Error bars show the standard deviation from three independent recordings with different pulled glass nanopores. The conductive current peaks were larger with increasingly negative voltages, while the dwell times shortened. **(E)** Event rate of the 20 nm diameter AuNPs translocation as a function of voltage. **(F)** Event rate of the 20 nm diameter AuNPs translocation as function of the sample concentration 6, 8, 12, and 25 µg/mL which correspond to concentrations of 1.4 x 10^9^, 1.1 x 10^10^, 7.1 x 10^10^, and 3.1 x 10^10^ particles/mL (calculated assuming all particles have the same average diameter). Error bars show the standard deviation from three independent recordings with different pulled glass nanopores.

### **S7: Nanopore ion current traces for AuNPs of different diameters.**

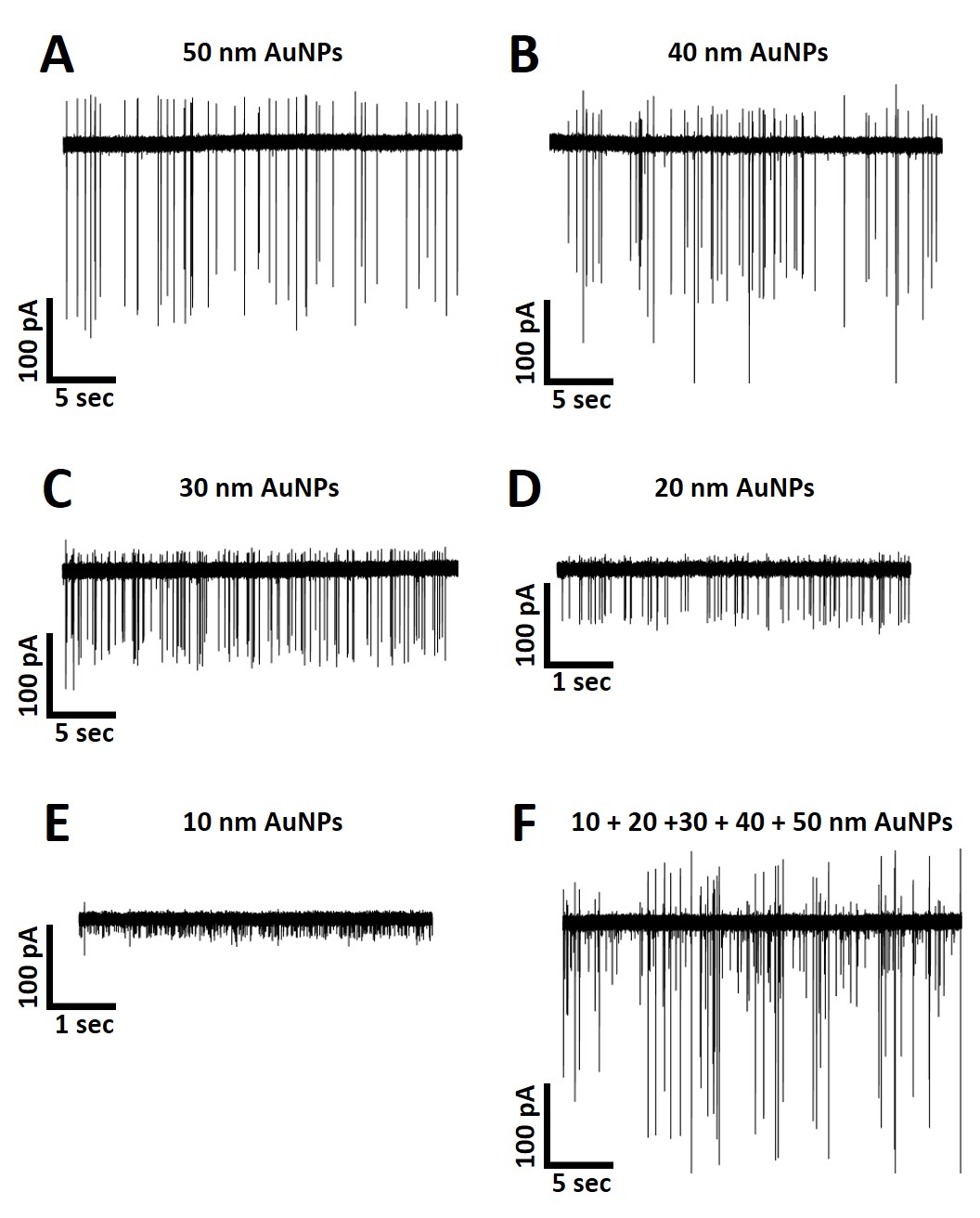

**Figure S7:** Ion current traces for PEG-carboxyl capped AuNP samples translocated through 60 nm diameter nanopores into an electrolyte bath of 50% PEG in 50 mM KCl. The pipettes were biased at -500 mV and filled with 50 mM KCl containing **(A)** 50 nm diameter AuNPs; **(B)** 40 nm diameter AuNPs; **(C)** 30 nm diameter AuNPs; **(D)** 20 nm diameter AuNPs; **(E)** 10 nm diameter AuNPs; or **(F)** or a mixture containing: 10, 20, 30, 40, and 50 nm AuNPs. All the ion current traces have the same current scale; however, parts D and E have a shorter timescale to allow showing the higher translocation frequency. The nanoparticles concentrations in panels (A)-(E) were 25 µg/mL, which corresponds to 3.0 x 10^11^, 3.1 x 10^10^, 1.1 x 10^10^, 3.5 x 10^10^, 2.1 x 10^9^ of particles/mL, respectively. The concentration of each component in panel F mixture was: 7.4 x 10^9^ of particles/mL for 10 nm AuNPs, 7.6 x 10^8^ of particles/mL for 20 nm AuNPs, 2.6 x 10^9^ of particles/mL for 30 nm AuNPs, 3.5 x 10^9^ of particles/mL for 40 nm AuNPs, 1.3 x 10^9^ of particles/mL for 50 nm AuNPs.

**Table S1:** Average conductive (ΔI_C_) and resistive (ΔI_R_) peak current values with the standard error (SE) obtained for PEG-carboxyl capped AuNP samples translocating through ~60 nm diameter nanopores into an electrolyte bath containing 50% PEG and 50 mM KCl. The pipette was biased at ‑500 mV, and filled with a solution containing 50 mM KCl and 25 µg/mL of nanoparticles. Note: we were unable to measure resistive peaks for the 10 and 20 nm diameter particles under these conditions. Each value shows the results based on a single nanopore recording with number of translocated events displayed in the table.

| **Pore diameter (nm)** | **NP diameter (nm)** | **NP diameter / pore diameter** | **Translocated events** | **ΔI_C_ ± SE**  **(pA)** | **ΔI_R_ ± SE**  **(pA)** |
| --- | --- | --- | --- | --- | --- |
| 60 | 50 | 0.83 | 286 | 225 ± 1 | 71 ± 1 |
|  | 40 | 0.67 | 469 | 160 ± 2 | 37 ± 1 |
|  | 30 | 0.50 | 937 | 98 ± 1 | 24 ± 1 |
|  | 20 | 0.33 | 1132 | 61 ± 1 | - |
|  | 10 | 0.17 | 1715 | 21 ± 1 | - |

### **S8: Biphasic signal reproducibility.**

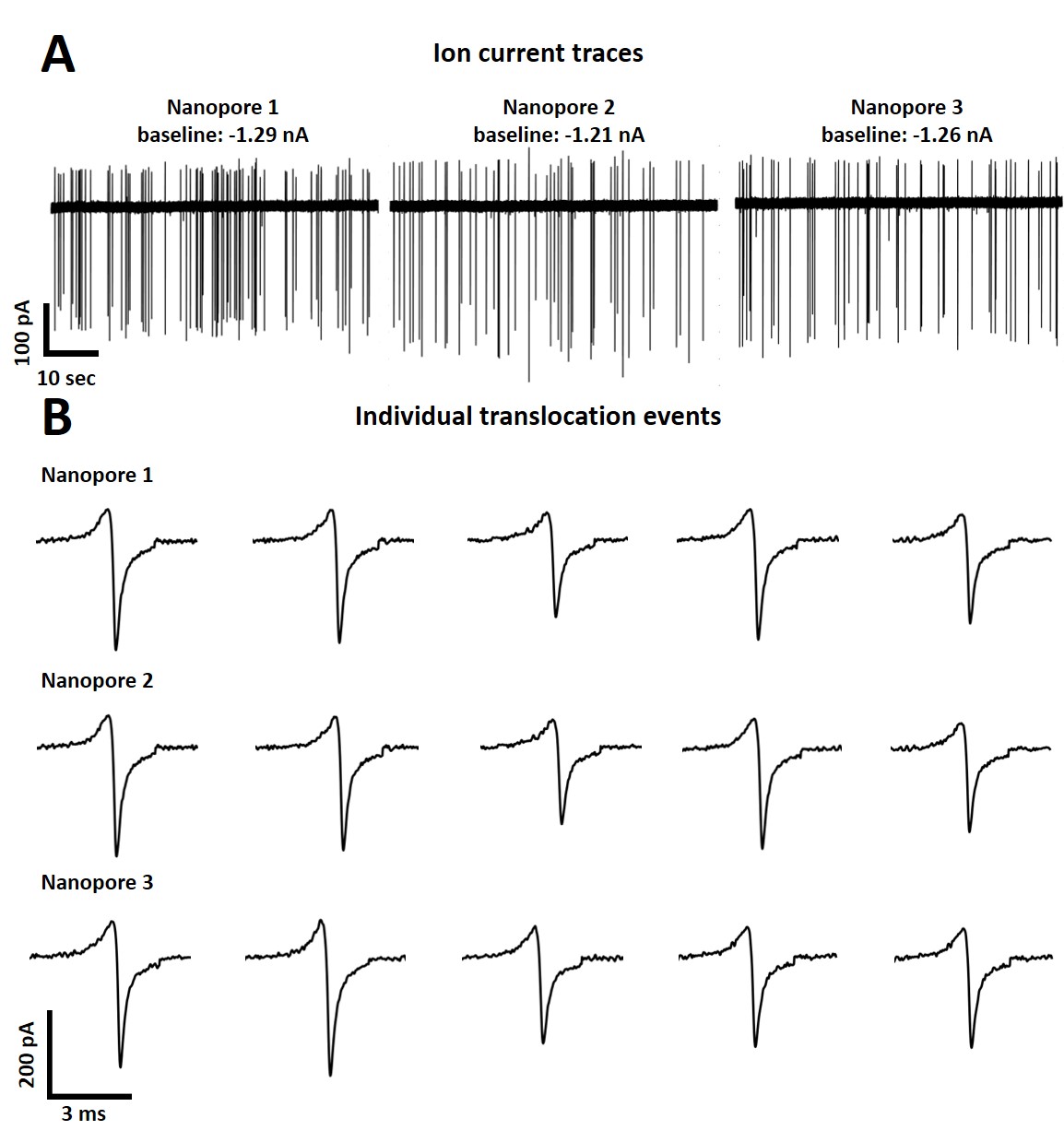

**Figure S8:** Biphasic signal reproducibility. **(A)** Ion current traces obtained with three different glass nanopores. All the ion current traces were adjusted to the same current and time scale. **(B)** Consecutive biphasic translocation events extracted from ion current traces recorded in (A) with three different glass nanopores. All the individual translocation events have the same current and time scales. Nanopore recordings were carried out using 60 nm diameter pore sizes and 50 nm diameter AuNPs translocated in 50% PEG and 50mM KCl and biased at -500 mV. A reproducible effect of the biphasic signal can be observed for each recording.

### **S9: Effect of the nanoparticle-nanopore size ratio on translocation signatures.**

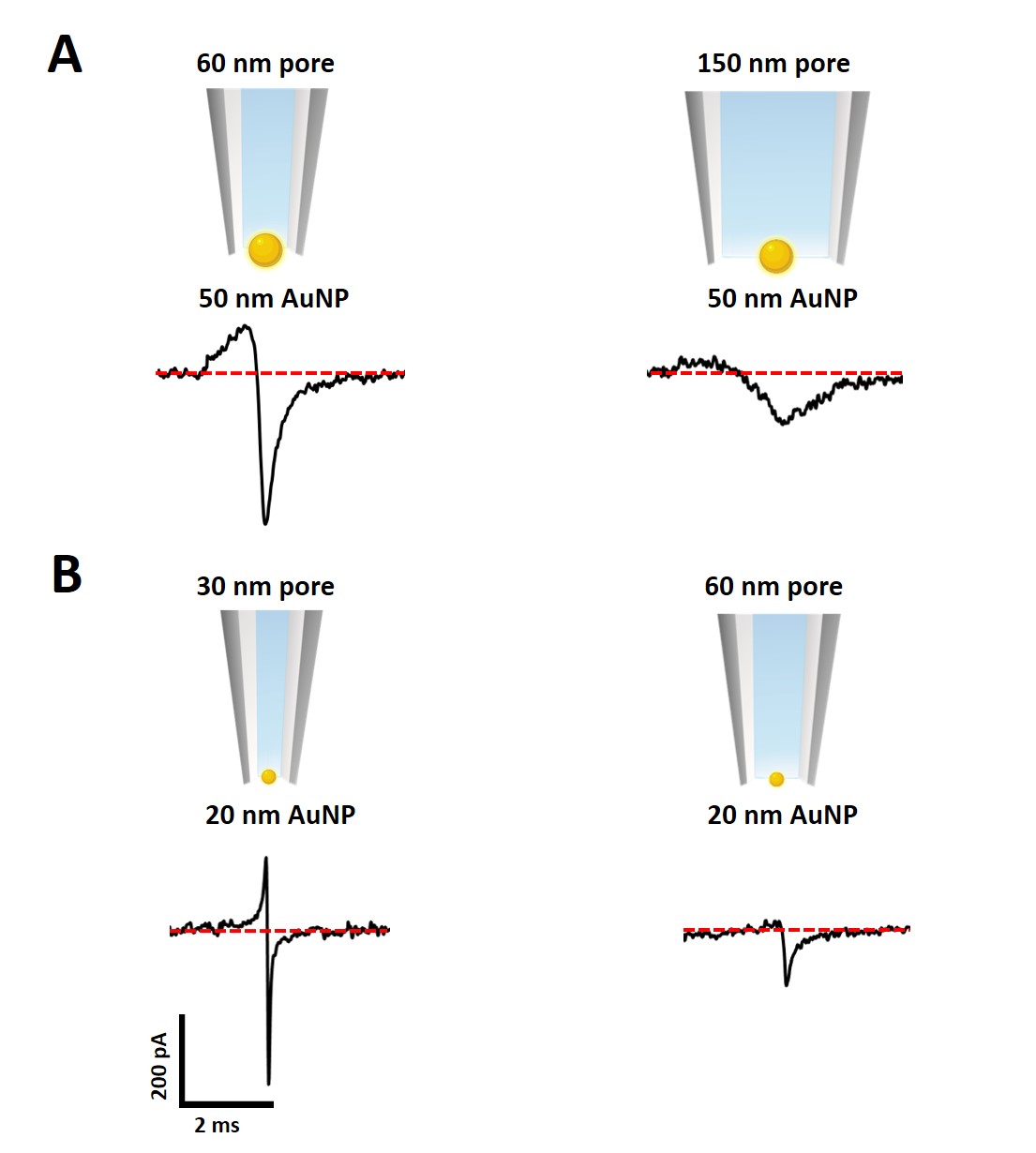

**Figure S9: (A)** Representative individual translocation peaks for 50 nm diameter AuNPs translocating through a 60 nm diameter pore (left) and 150 nm diameter pore diameter (right), showing the transition from biphasic to conductive shape when the pore diameter increases with respect to the diameter of the particle. **(B)** Representative individual translocation peaks for the 20 diameter nm AuNP translocated through a 30 diameter nm pore (left) and 60 nm pore diameter (right), showing the transition from biphasic to conductive peak shape when the pore diameter is increased with respect to the diameter of the particle. The current and time scales are the same for all the translocation peaks. Translocation was carried out in 50% PEG in 50 mM KCl and the pipettes were filled with 50 mM KCl and 25 µg/mL nanoparticles and biased at -500 mV.

### **S10: Characterization of metallic nanoparticles.**

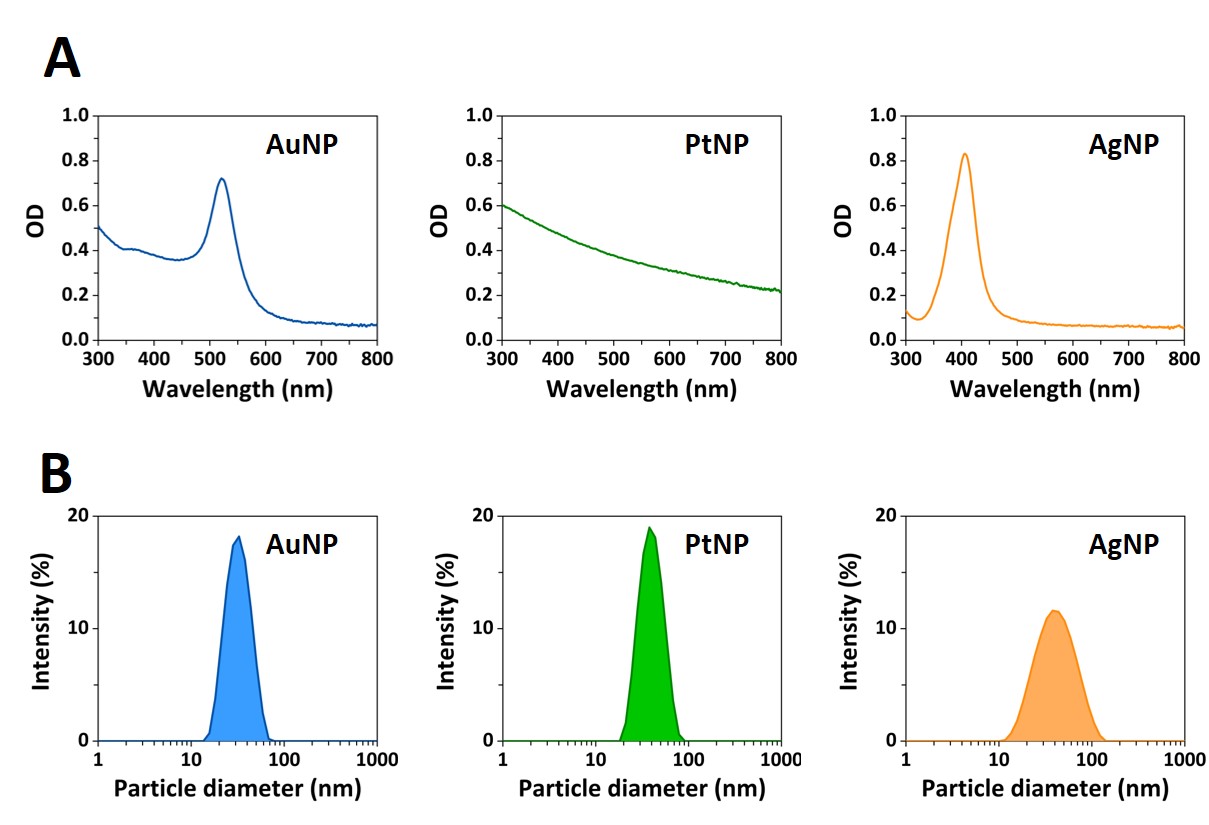

**Figure S10:** Characterization of the homogeneous metallic nanosphere samples. **(A)** UV-Vis spectra of nominally 30 nm diameter nanospheres in 10 mM KCl: PEG carboxyl-capped AuNPs (left panel), citrate- capped PtNPs (middle panel), citrate-capped AgNPs (right panel). A strong plasmon resonance optical absorbance at 520 nm was recorded for the AuNPs (left panel) and 420 nm for the AgNPs (right panel). No plasmon resonance was observed for the PtNPs (middle panel). **(B)** Dynamic light scattering (DLS) measured size distribution (hydrodynamic diameter) of nominally 30 nm diameter nanospheres in 10 mM KCl: PEG carboxyl-capped AuNPs (left panel), citrate- capped PtNPs (middle panel), citrate-capped AgNPs (right panel). The size distribution measured shows the presence of a homogenous suspension of nanospheres without aggregates formation.

### **S11: Nanopore translocation AuNS samples.**

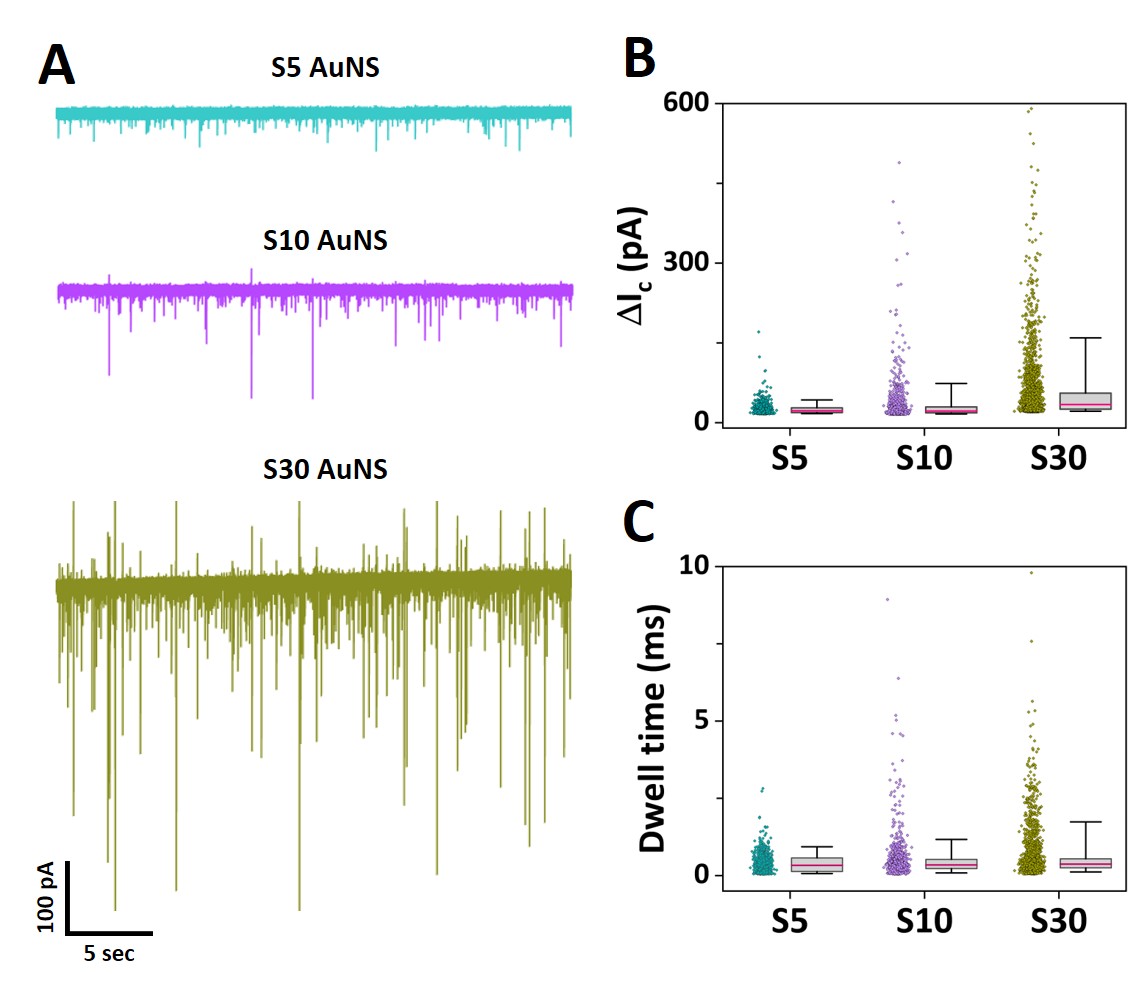

**Figure S11: (A)** Current-time traces obtained for S5, S10, and S30 gold nanostars in 50% PEG in 10 mM KCl conditions using a fixed pore size (80 nm diameter) biased at -700 mV. The nanoparticle concentrations for S5-S30 correspond to: 6.5 x 10^9^, 5.0 x 10^11^, and 3.9 x 10^11^ particles/mL. The current and time scales are the same for all the ion current traces. (S5 AuNS: 751 events; S10 AuNS: 1115 events; S30 AuNS: 2589). Box and whisker plots showing the conductive peak current **(B)** and the dwell time **(C)** for the S5, S10, and S30 AuNS samples (median, interquartile range 25-75%, and whiskers range 5-95%).

### **S12: AuNS samples characterization.**

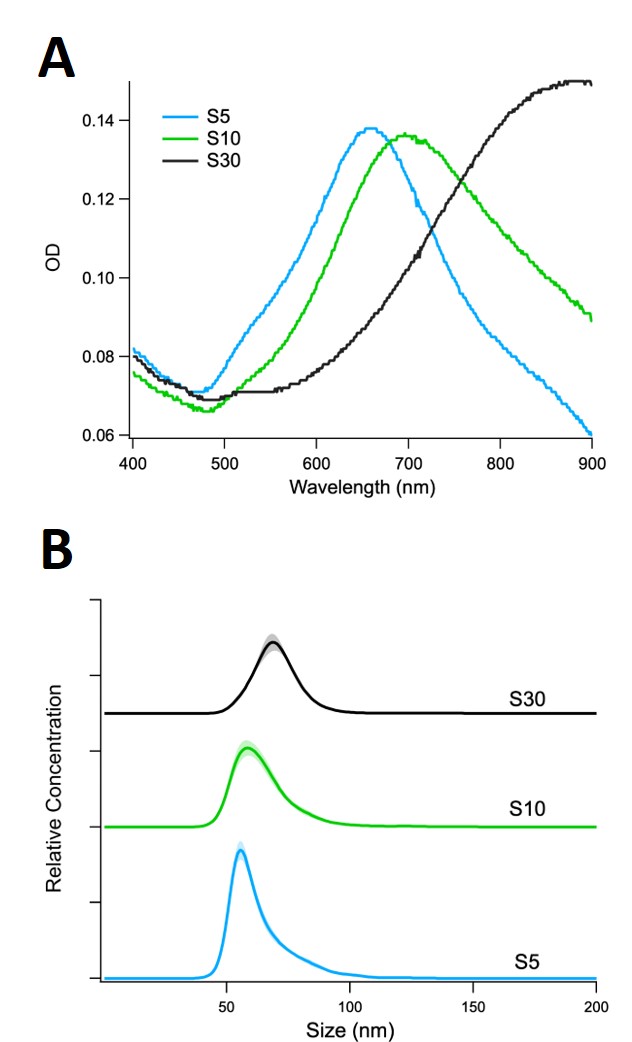

**Figure S12:** Characterization of the synthesized gold nanostar samples prepared in Mili-Q water. **(A)** Extinction spectra of the synthesized gold nanostars samples: S5 (blue line: ʎ_max_ = 658 nm), S10 (green line: ʎ_max_ = 696 nm), and S30 (black line: ʎ_max_ = 888 nm). A shift to higher wavelength is observed with increasing density of the nanostar branches. **(B)** Nanoparticle tracking analysis spectra of the gold nanostar samples: S5 (blue line), S10 (green line), and S30 (black line). The mean size (hydrodynamic diameter) determined was: 63.4 ± 0.3 for S5 sample, 65.4 ± 0.3 for S10 sample, and 70.7.4 ± 0.3 for S30 sample. The multiple AuNS morphologies present (see Figure S13) are responsible for the skewed and broad hydrodynamic size distribution.

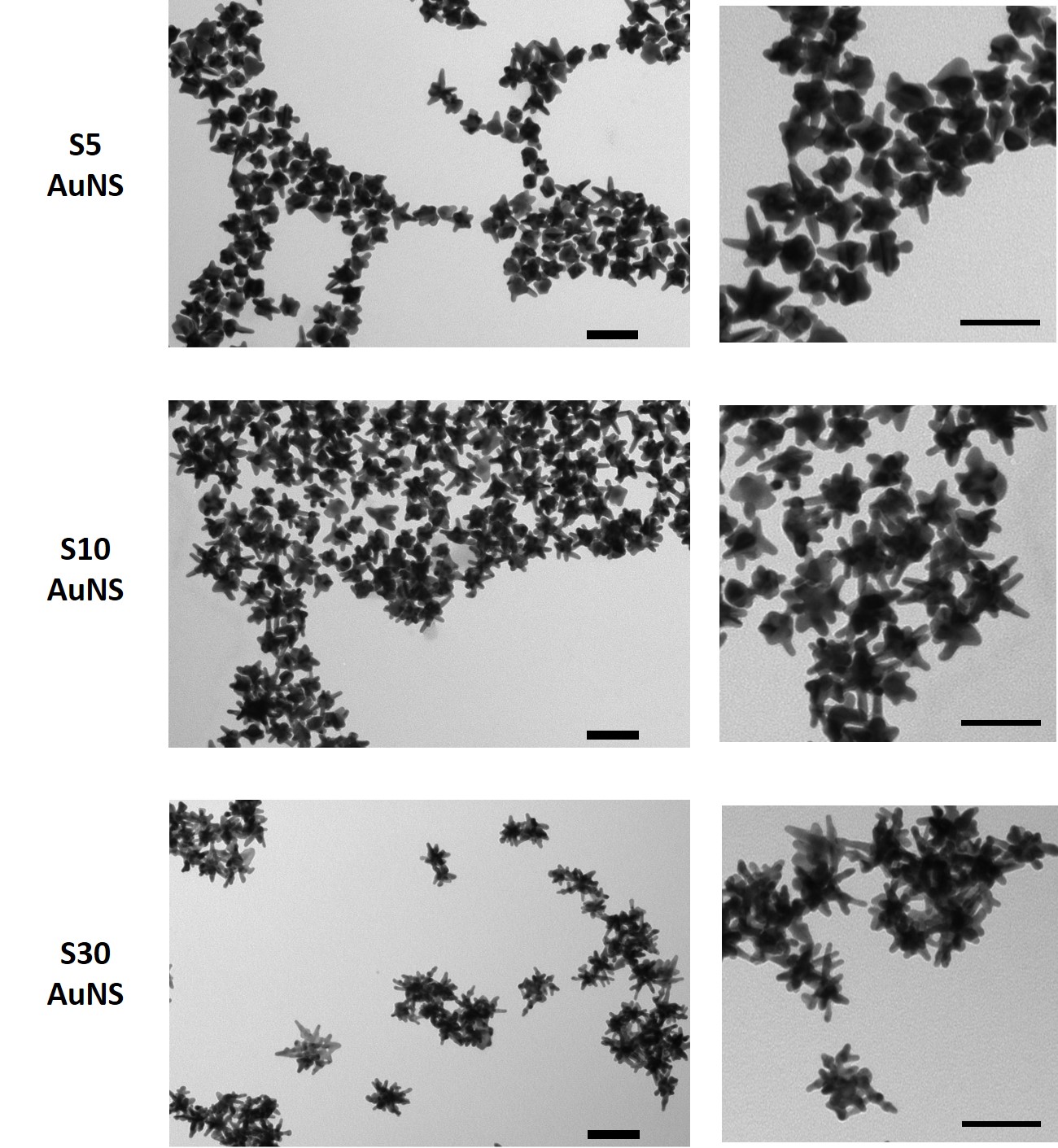

**Figure S13:** TEM images of the synthesized gold nanostars: S5 (top panel), S10 (middle panel); S30 (bottom panel). Scale bar graphs: 100 nm.

### **S13: β-gal SNAs synthesis and characterization.**

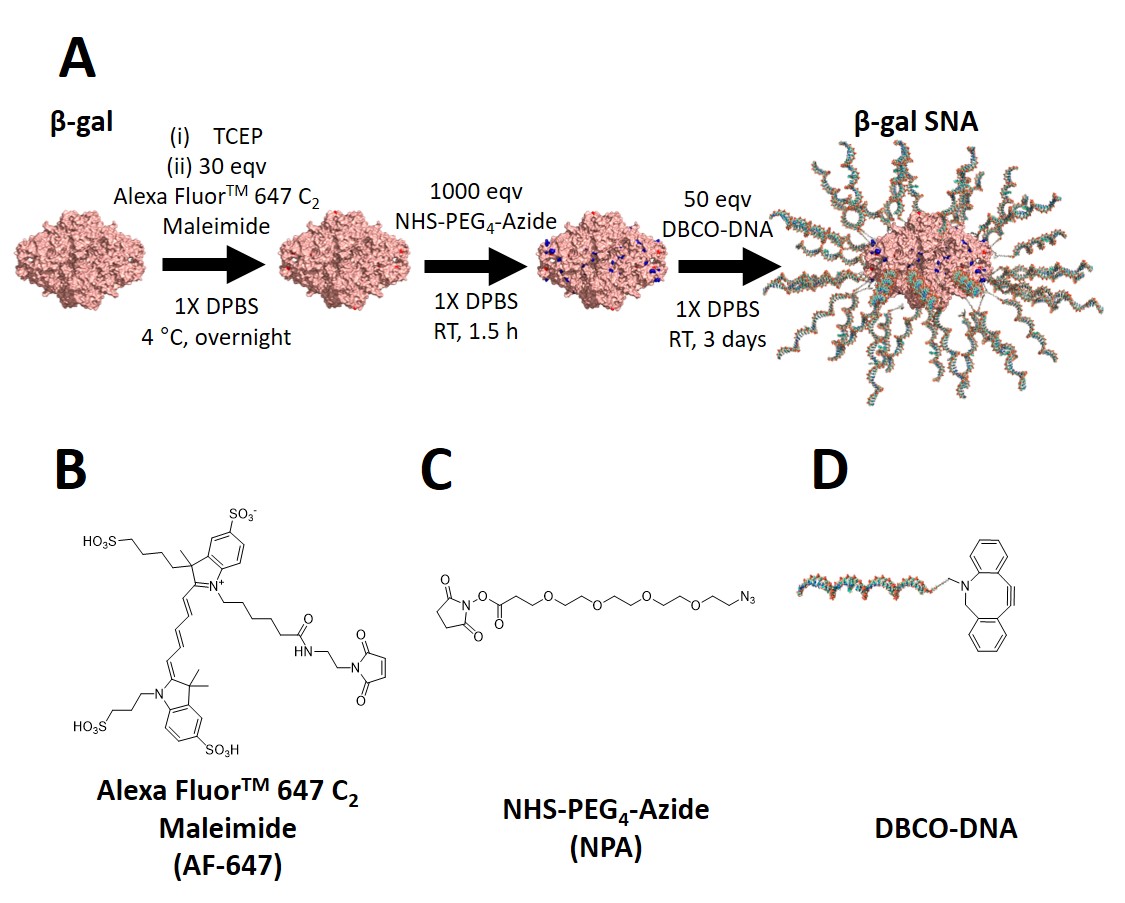

**Figure S14.** **(A)** Schematic depicting the synthesis steps of β-gal SNAs. Structures of **(B)** AF-647, **(C)** NHS‑PEG4-azide, and **(D)** DBCO-DNA.

Figure S14 shows the synthesis steps of obtaining β-gal SNAs. A stock solution of β-gal (Sigma, Cat. No. 10105031001) was exchanged into DPBS (Sigma, Cat. No. D8537) through centrifuge filtration (15000 rpm, 3 min, 4 °C), three times washing using a centrifuge (Thermo Fisher, Cat. No. 5425) using 100 kDa MWCO Amicon filters (Thermo Fisher, Cat. No. UFC510096). The β-gal was then treated with TCEP (4 nM final concentration) to reduce the disulphide bonds and vortexed. Thereafter, the solution was passed through a NAP-5 column to remove the TCEP and the purified β-gal solution was concentrated using a 100 kDa MWCO Amicon filter. The protein concentration was determined via UV-vis spectroscopy using the extinction coefficient at the characteristic protein peak at 280 nm. Then 30-fold molar excess of Alexa Fluor 647 C_2_ Maleimide (AF-647, Thermo Fisher, Invitrogen, Cat. No. A20347) was added to the protein and the mixture was shaken overnight (1000 rpm, 4 °C) using a Benchmark MultiTherm Heating and Cooling Shaker (Marshall Scientific, Cat. No. H5000-HC). The solution was then rinsed multiple times with 0.1 M NaHCO_3_ (Thermo Fisher, Cat. No. BP328-500) using a 100 kDa MWCO Amicon filter to remove any unreacted dye. The absorbance of the filtrate was monitored at 650 nm to track the free dye. The washing was stopped when the absorbance at 650 nm was negligible. Using UV-vis spectroscopy, the concentration of the protein ($c_{\beta-gal}$) and the number of dyes attached per protein ($N_{dye}$) were calculated using equations SE1 and SE2:

$c_{\beta-gal}=\frac{A_{280 nm}-\varepsilon_{280 nm}^{AF-647}c_{AF-647}}{\varepsilon_{280 nm}^{\beta-Gal}}$	(SE1)

$N_{dye}=\frac{c_{AF-647}}{c_{\beta-gal}}$	(SE2)

$c_{AF-647}$ is first determined using Equation S1, where $\varepsilon_{650 nm}^{AF-647}$ = 270,000 M^-1^ cm^-1^. The value for $\varepsilon_{280 nm}^{AF-647}$ is 8,100 M^-1^ cm^-1^.

Thereafter, NHS-PEG_4_-azide (Thermo Scientific, Cat. No. PI26130) was added to the dye-labelled β-gal (β-gal-AF) at 1000-fold molar excess and the reaction was incubated for 1.5 hours with shaking (1000 rpm, RT). The reaction mixture was then run through a NAP™-25 column (Cytiva illustra, Cat. No. 17085202) and washed 5 times using centrifuge filtration (100 kDa MWCO Amicon filters, 15000 rpm, 4 °C, 3 min per wash) to separate any free azide molecules. The protein concentration of the product (β-gal-AF-azide) was again determined via UV-vis spectroscopy. Then 50-fold molar excess of DBCO-DNA was added to β-gal-AF-azide (samples were prepared with both DBCO-DNA 1 and DBCO-DNA 2; see Table S2 for sequences). The reaction was shaken for 3 days (1000 rpm, RT), following which the reaction mixture was washed several times using PBS and 100 kDa MWCO Amicon filters (15000 rpm, 4 °C, 6 min per wash) to remove free DNA. The final product (β-gal SNAs) was characterized by UV-vis spectroscopy. First, based on the absorbance at 647 nm, $c_{AF-647}$ was first determined. Then, using SE2, $c_{\beta-gal}$ was determined. Then, using the absorbance at 260 nm, the number of DNA per protein ($N_{DNA}$) was determined using equation SE3:

$N_{DNA}=\frac{A_{260 nm}-\varepsilon_{260 nm}^{\beta-Gal} c_{\beta-gal}}{\varepsilon_{260 nm}^{DNA}c_{\beta-gal}}$	(SE3)

Based on these calculations, we determined that the final β-gal SNAs employing DBCO-DNA 1 had an average of ~2 dyes and ~42 DNA strands on the protein surface whereas those using DBCO-DNA 2 had ~5 dyes and ~22 DNA strands on the protein surface. Figure S15 shows UV-Vis spectra of β-gal derivates. The absorbance spectrum of native β-gal shows a peak at 280 nm, however, once β-gal is modified with AF-647 (β-gal-AF), an additional absorbance peak is observed at 647 nm corresponding to AF-647. β-gal SNAs show absorbance peaks at both 260 nm (corresponding to DNA) and 647 nm (corresponding to the AF-647). The number of dyes and DNA strands per protein calculated using the absorbance spectra below indicate that on an average, ~2 dyes and ~42 DNA strands on the protein surface for DBCO-DNA 1 and ~5 dyes and ~22 DNA strands on the protein surface for DBCO-DNA 2. Representative data is shown for β-gal SNA_22_ with DBCO-DNA 2 (Figure S15).

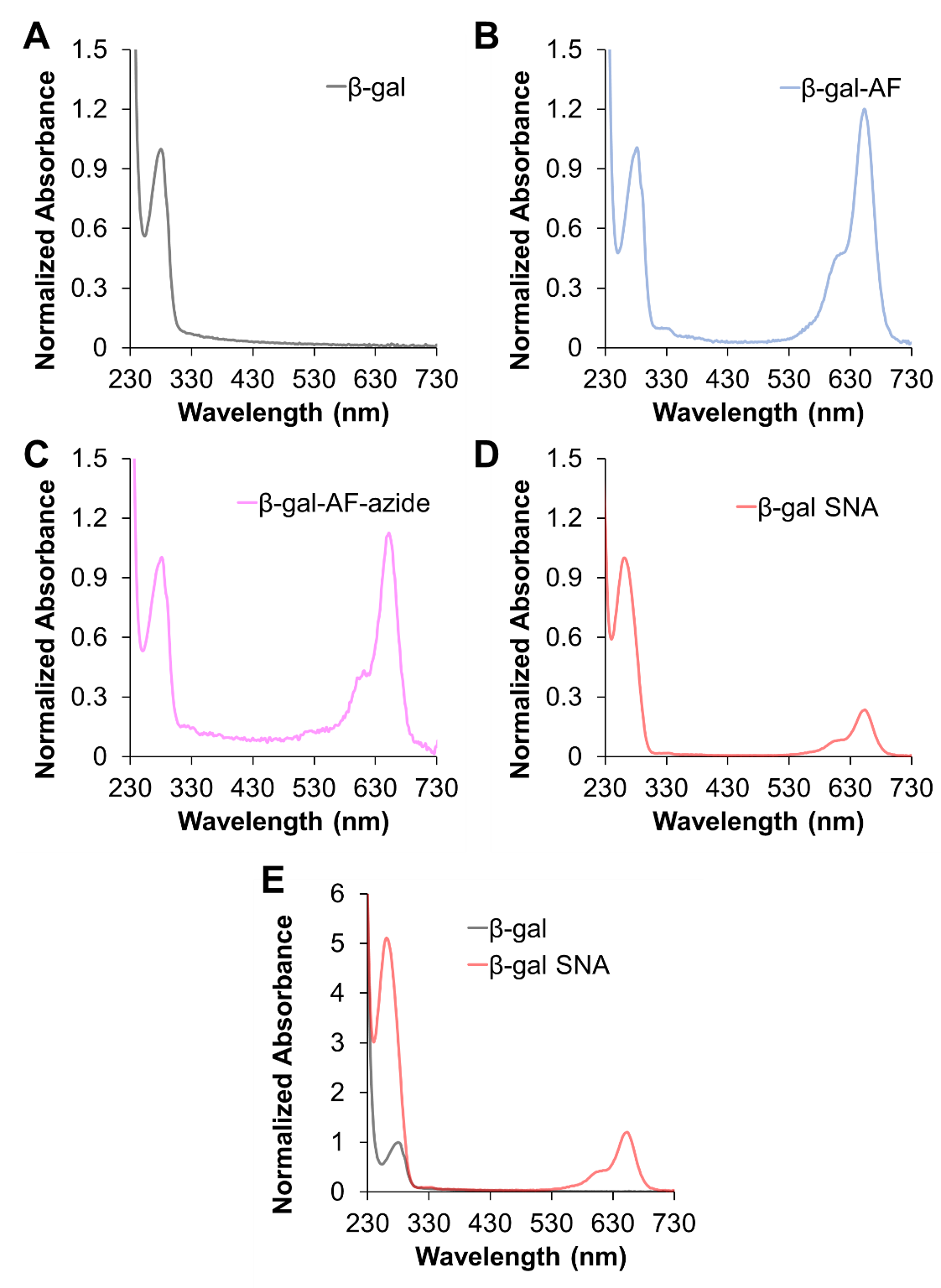

**Figure S15.** Representative UV-Vis spectra of **(A)** native β-gal, **(B)** β-gal-AF, **(C)** β-gal-AF-azide, and **(D)** β-gal SNA_22_. **(E)** Absorbance of identical concentrations of β-gal and β-gal in Mili-Q water. In all graphs, the absorbance is normalized to the absorbance at 280 nm (corresponding to the characteristic protein absorbance) is normalized to a value of 1.

Native gel electrophoresis was used to characterize β-gal SNAs. For the PAGE gel electrophoresis, 10 µL of 1 µM SNAs (by protein) was added to 2 µL of glycerol and loaded onto a 4-15% acrylamide gel (Bio-Rad, 4-15% Mini-PROTEAN TGX Precast Protein Gels, 10-well, Cat. No. 4561084) and run for 60 min at 100V at room temperature. The running buffer was prepared by diluting 10X Tris-Borate-EDTA (TBE Buffer, Thermo Fisher, Cat. No. BP13334) to obtain a 1X TBE Buffer. The native protein was also run as a control. 5 µL EZ-Run Prestained Rec Protein Ladder (Thermo Fisher, Cat. No. BP3603500) was also used. For β-gal, after electrophoresis, a solution of 10 mM ortho-nitrophenyl-β-D-galactopyranoside (ONPG, Sigma, Cat. No. N1127) the protein substrate was added to the gel to identify the positions of active proteins. Thereafter, the gels were stained with GelRed Nucleic Acid Gel Stain following the manufacturer's protocol and imaged using the ChemiDoc MP Imaging System (Bio-Rad, USA). Subsequently, the gels were stained with SimplyBlue SafeStain (Thermo Fisher, Cat. No. LC6065) following the manufacturer's protocol and reimaged. Both β-gal and β-gal SNA migrate efficiently through the PAGE gel (as the isoelectric point is < 7). When the gel is treated with 10 mM ONPG (ortho-nitrophenyl-β-D-galactopyranoside), a substrate of β-gal, a yellow color is observed due to the formation of o-nitrophenol (Figure S16A). This further confirms that the bands contain active β-gal protein. The position of the β-gal with respect to the protein ladder suggest that we are looking at one subunit of the tetramer. Figure S16B-D show that β-gal SNA_22_ how signal in the AF-647, GelRed, and Coomassie channels, indicating the SNAs are comprised of dye, DNA, and protein, respectively. In contrast, native β-gal is only visible in the Coomassie channel as Coomassie stains the protein.

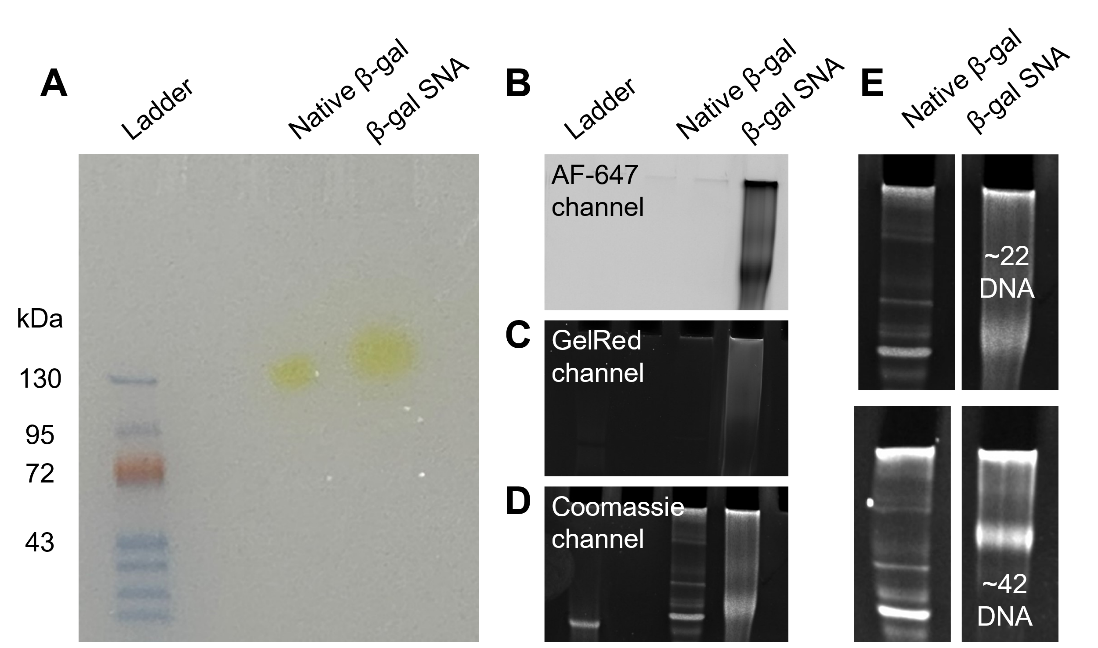

**Figure S16.** Native PAGE images of β-gal and β-gal SNA_22_. **(A)** Photograph of the native gel after exposing the gel to ONPG substrate solution. β-gal results in the conversion of ONPG to a yellow-colored product, o-nitrophenol. **(B)** Native gel imaged in the AF-647 channel. The β-gal SNA_44_ is visible as it has the AF-647 modification, but the native β-gal is not. **(C)** Native gel imaged after staining with GelRed Nucleic Acid Gel Stain. The native β-gal is not visible whereas β-gal SNA_44_ is visible (the band appears darker, likely due to high DNA concentration). **(D)** Native gel imaged after Coomassie protein staining using SimplyBlue SafeStain.

**Table S2.** DNA sequences used for ProSNA samples.

| Sequence name | Sequence (5’-3’) | ε_260_  (M^-1^cm^-1^) | Expected mass  (m/z) | Observed mass  (m/z) |
| --- | --- | --- | --- | --- |
| DBCO-DNA 1 | DBCO TEG-TTTT AAG ACG AAT ATT TAA GAA | 240,800 | 7351 | 7250 |
| DBCO-DNA 2 | DBCO TEG-TTTT AAC GAC TCA TAT TAA CAA | 228,600 | 7247 | 7162 |
| Thiol-DNA | C6-SS-AAG ACG AAT ATT TAA GAA | 200,500 | 5760 | 5787 |

C6-SS: Thiol-Modifier C6 S-S (1-O-Dimethoxytrityl-hexyl-disulfide,1’-[(2-cyanoethyl)-(N,N-diisopropyl)]-phosphoramidite)

DBCO-TEG: 5’-DBCO-TEG Phosphoramidite (10-(6-oxo-6-(dibenzo[b,f]azacyclooct-4-yn-1-yl)-capramido-N-ethyl)-O-triethyleneglycol-1-[(2-cyanoethyl)-(N,N-diisopropyl)]-phosphoramidite)

### **S14: Translocation events of β-gal SNAs.**

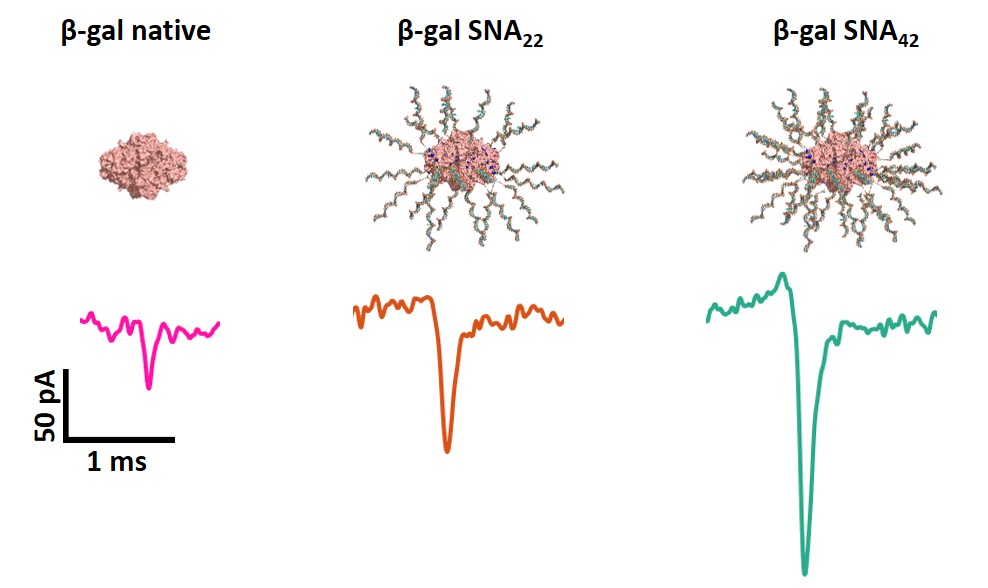

**Figure S17.** Representative individual translocation events recorded for ProSNA samples: β-gal native protein, β-gal SNA_22_, and β-gal SNA_42_ from a 30 nm pore diameter biased at -700 mV filled with 100 mM KCl and 5 nM β-gal derivatives into a bath containing 50% PEG in 100 mM KCl. The current and time scales are the same for all the translocation peaks.

**Table S3.** Average conductive peak current (ΔI_C_) and dwell time values with their standard error (SE) obtained for the ProSNA samples translocated through a 30 nm nanopore diameter biased at -700 mV. The nanopore was filled with 100 mM KCl and 5 nM β-gal derivatives, whereas the external bath contained 50% PEG and 100 mM KCl.

| **ProSNA sample** | **Translocated events** | **ΔI_C_ ± SE**  **(pA)** | **Dwell time ± SE**  **(ms)** |
| --- | --- | --- | --- |
| β-gal NT | 262 | 32 ± 1 | 0.10 ± 0.01 |
| β-gal SNA_22_ | 2042 | 94 ± 1 | 0.11 ± 0.01 |
| β-gal SNA_42_ | 1224 | 171 ± 1 | 0.13 ± 0.01 |

### **S15: Finite element modelling.**

To provide a mechanistic understanding of the experimentally observed current responses during nanoparticle translocation, we developed numerical simulations describing the electric potential, $\phi$, ion concentrations, $c_{i}$, ($i$ = K^+^ or Cl^-^), fluid flow, $\boldsymbol{u,}$and pressure, $p$, in the region around the aperture of the glass nanopore as shown in Figure S18. These simulations, which are based on our previous work describing a nanopore immersed in a polymer electrolyte (2), are described in detail below.

The equations were solved using the commercial finite element software COMSOL Multiphysics (version 5.6). To aid reproduction of the model, a COMSOL ‘Model Report’ containing all the parameter values, equations solved, boundary conditions and simulation settings is included as a separate Supporting Information file.

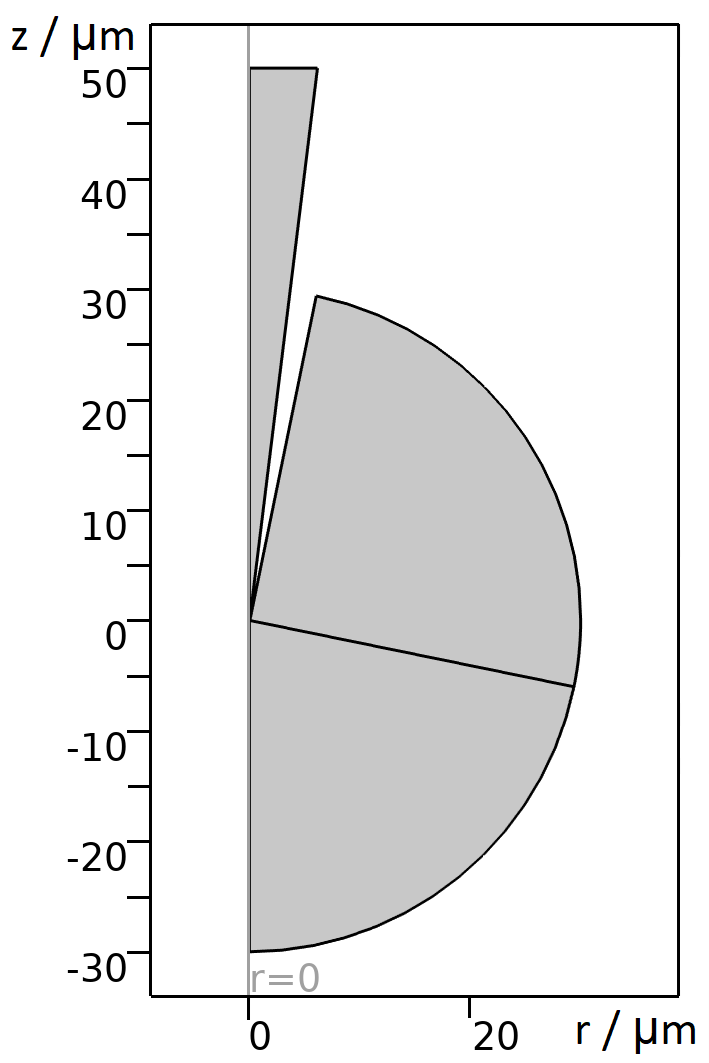

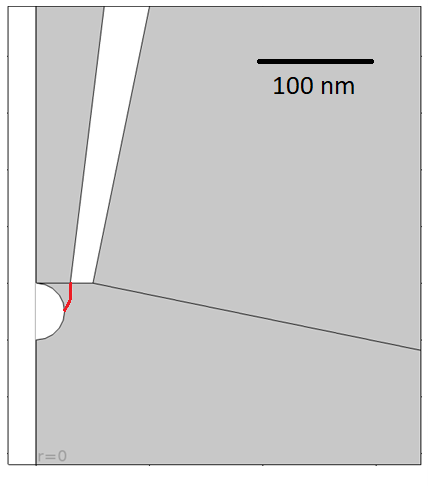

**Figure S18.** Geometry used for the numerical simulations. Left-hand side shows the full domain of simulation while the right-hand side shows a zoomed-in view close to the end of the pipette where a nanoparticle has just exited the glass nanopore. The red line is the interface between the internal (KCl only) solution and the external solution (PEG + KCl). Note that the interface shape changed according to the position of the nanoparticle along the z-axis.

**Equations for concentration, electric potential, and fluid flow**

The ion concentrations were determined by solving the continuity equation (SE4) where the ionic flux ${\overset{⃑}{J}}_{i}$ is given by the Nernst-Planck equation (equation SE5):

|  | $\frac{\partial c_{i}}{\partial t}=-\nabla\cdot{\overset{⃑}{J}}_{i}$ | (SE4) |
| --- | --- | --- |
|  | ${\overset{⃑}{J}}_{i}=-D_{i}^{\alpha}\nabla c_{i}+\frac{z_{i}F}{RT}D_{i}^{\alpha}c_{i}\nabla\phi+\boldsymbol{u}c_{i}$ | (SE5) |

in which $D_{i}^{\alpha}$ is the diffusion coefficient of ion $i$ in phase *α* (PEG+KCl/KCl), $z_{i}$ the ion charge, $F$ Faraday’s constant, $R$ the universal gas constant, and $T$ the temperature. The values for these and all other constants from the simulations are listed below in Table S4 and in the COMSOL model report file.

The electric potential is determined by solving Poisson’s equation (SE6):

| $\nabla^{2}\phi=-\frac{F}{\varepsilon_{0}\varepsilon_{r,\alpha}}\sum_{i} z_{i}c_{i}$ | (SE6), |
| --- | --- |

$\varepsilon_{0}$ is the permittivity of free space and $\varepsilon_{r,\alpha}$ is the relative dielectric constant of the electrolyte phase. The space‑charge term (right-hand side) comes from the ions in solution.

The fluid flow and pressure are determined by solving the Navier-Stokes equations for an incompressible fluid including an electroosmotic volume force contribution (SE7a,b)

|  | $\boldsymbol{(u}\cdot\nabla)\boldsymbol{u}=\frac{1}{\rho_{\alpha}}\left( -\nabla p+\eta_{\alpha}\nabla^{2}\boldsymbol{u}-F\left( \sum_{i} z_{i}c_{i} \right)\nabla\phi\right)$ | (SE7a) |
| --- | --- | --- |
|  | $\rho_{\alpha}\nabla\cdot\boldsymbol{u}=0$ | (SE7b) |

where $\rho_{\alpha}$ is the solution density in phase $\alpha$, and $\eta_{\alpha}$ the dynamic viscosity.

**Boundary conditions**

*Electrostatics*

The wall of the pipette holds a negative charge density, *σ*_glass_, while the nanoparticle had a surface charge, *σ*_NP_, which is encoded in the model with the boundary condition:

$$\hat{n}\cdot\nabla\phi=-\frac{\sigma_{x}}{\varepsilon_{0}\varepsilon_{r,\alpha}}$$

where $x$ = glass or NP.

The surface charge of the glass nanopore wall, $\sigma_{glass}$, was selected according to the parametric study performed in a previously developed model (2).

The surface charge of a 50 nm Au nanoparticle, $\sigma_{np}$, was calculated analytically based on the following equation (3) which describes the Gouy-Chapman equation for a flat charged surface and relates zeta potential $\left( \zeta\right)$ and effective charge density (sum of charge density of flat surface and its surrounding ions in the slipping plane):

$$\sigma_{eff,NP}=\sqrt{8c_{avg}N_{A}\varepsilon_{r}\varepsilon_{0}k_{B}T}\sinh\left( \frac{q_{e}\zeta}{k_{B}T} \right)$$

where $c_{avg}$: concentration of solution (2 mM citrate in zeta potential measurements), $N_{A}$: Avogadro’s number, $\varepsilon_{r}=77$: relative electrical permittivity of solution, $k_{B}$: Boltzmann’s constant and $q_{e}$: elementary charge.

In the external solution, we set the potential to *ϕ* = 0 V, while the internal solution is set at *ϕ* = *V*_app_. The potential is corrected to match the Gouy-Chapman description of the electric double layer where the boundaries reach the glass.

*Concentration*

The concentration at the interior and exterior solution boundaries is chosen to match the bulk concentrations used in the experiments. As with the electrostatics, a correction is used within the electric double layer to match Gouy-Chapman theory. A no flux $(\hat{n}\cdot{\overset{⃑}{J}}_{i}$= 0) boundary condition is applied on the glass walls.

*Fluid flow*

A no-slip boundary condition ($\boldsymbol{u}=0)$ was used on the glass and nanoparticle walls. A pressure difference equal to the hydrostatic pressure (calculated from the pore height) was applied between the external solution boundary and that inside the pipette.

**Geometry**

The simulations all used the 2D axisymmetric geometry shown in Figure 3 where the particle resides on the axis of symmetry of the glass nanopore. The vertical height of the particle was varied to allow us to understand the current response. For simplicity, the interface between the interior (KCl) and bath (PEG + KCl) was assumed to have zero width. When the particle was inside the pipette (not shown), the interface was set to coincide with the aperture of the pipette, whereas when the particle was translocating outward, the particle distorts this interface (red line in Figure S18). As we do not have any experimental data on the geometry of this deformation, we approximate it by a circular arc between the equator of the particle and the pipette aperture with radius of curvature equal to the pore radius.

**Mesh**

A boundary layer mesh with a thickness equal to one tenth of the reciprocal Debye length was used on the surface of the nanoparticle and glass. Small mesh elements were used near interface between the PEG and the KCl solutions and the orifice of the pore to accurately capture the strong gradients in this region. See the COMSOL model support (addition Supplementary Information file) for the full specifics of the mesh.

**Calculation of current**

The current is calculated from the integration of the ionic fluxes across the top of the pipette ($\partial\Omega)$ as defined by:

| $i=2\pi rF\int_{\partial\Omega} \left( {\overset{⃑}{J}}_{K^{+}}-{\overset{⃑}{J}}_{{Cl}^{-}} \right)\cdot\hat{n} ds$ | (SE8) |
| --- | --- |

**Physical Parameters**

**Table S4.** Key physical parameters used in the simulations alongside notes on how they were obtained. A detailed list of all parameters can be found in the COMSOL model report file.

| **Parameter** | **Value** | **Notes** |
| --- | --- | --- |
| $D_{K^{+}}^{PEG}$ (m^2^/s) | $1.59*{10}^{-10}$ | *best fit to model* |
| $D_{Cl^{-}}^{PEG}$ (m^2^/s) | $2.96*{10}^{-10}$ | *best fit to model* |
| $D_{K^{+}}^{no PEG}$ (m^2^/s) | $1.6*{10}^{-9}$ | *(4)* |
| $D_{{Cl}^{-}}^{no PEG}$ (m^2^/s) | $1.7*{10}^{-9}$ | *(4)* |
| $\kappa^{PEG}$ (S/m) | $0.613$ | *experimental* |
| $\kappa^{no PEG}$ (S/m) | 0.0855 | *experimental* |
| $\eta_{PEG}$ (Pa*s) | 8.73 | *experimental* |
| $\eta_{no PEG}$ (Pa*s) | $8.9 * {10}^{-4}$ | *for 0.059 (mol/kg) (5)* |
| $\rho_{PEG}$ (kg/m^3^) | 999.9 | *for 0.059 (mol/kg) (5)* |
| $\rho_{no PEG}$ (kg/m^3^) | 1086.2 | *for 50% PEG 8000 in H_2_O (kg/m^3^) (6)* |
| $c_{avg}$ (mM) | 50 | *experimental* |
| $\sigma_{glass}$ (mC/m^2^) | -12 | *best fit to model (7)* |
| $\sigma_{NP}$ (mC/m^2^) | -5.89 | *analytical calculation for 50 nm Au NP (3)* |

The values labelled as “experimental” were measured experimentally while the ones labelled as “best fit to model” were selected so that the simulated current where no nanoparticle was included in the geometry ($i_{0,sim}=-1.2983 nA$) was in good agreement with the experimentally obtained baseline current $\left( i_{0,exp}=-1.2 nA \right)$ at $V_{app}=-500 mV$ for the above conditions.

### **S16: Simulated nanoparticle surface charge effect.**

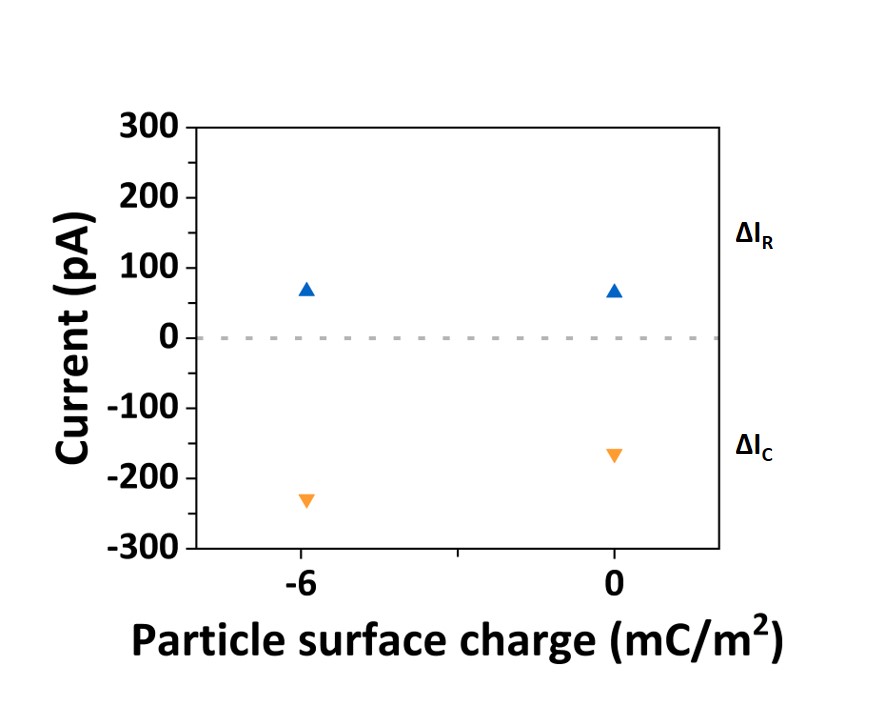

**Figure S19.** Simulated nanoparticle surface charge effect on the peak current magnitude for a 50 nm (negatively charged and neutral) particle translocating through a 60 nm pore in 50% PEG and 50 mM KCl conditions at -500 mV. Similar magnitudes of the resistive peak current are obtained in the case of negatively charged and neutral nanoparticle (blue points). The conductive peak current difference between the negatively charged and neutral particle denotes the effect of the external polymer electrolyte interface (50% PEG in 50 mM KCl) on the enhancement of the conductive peak current (orange points). The current scale denotes the ΔI with negative values representative of the conductive peak current contribution (ΔI_C_) relative to the current baseline (dotted grey line) and the positive values representative of the resistive peak current contribution (ΔI_R_) relative to the current baseline.

### **References**

1.	Plesa, C., and C. Dekker. 2015. Data analysis methods for solid-state nanopores. *Nanotechnology*. 26(8):084003

2.	Marcuccio, F., D. Soulias, C. C. C. Chau, S. E. Radford, E. Hewitt, P. Actis, and M. A. Edwards. 2023. Mechanistic Study of the Conductance and Enhanced Single-Molecule Detection in a Polymer–Electrolyte Nanopore. *ACS Nanoscience Au*. doi: 10.1021/acsnanoscienceau.2c00050.

3.	Ge, Z., and Y. Wang. 2017. Estimation of Nanodiamond Surface Charge Density from Zeta Potential and Molecular Dynamics Simulations. *The Journal of Physical Chemistry B*. 121(15):3394-3402, doi: 10.1021/acs.jpcb.6b08589.

4.	Haynes, W. M. 2016. CRC handbook of chemistry and physics 97th Edition. CRC press.

5.	Hai-Lang, Z., and H. Shi-Jun. 1996. Viscosity and density of water+ sodium chloride+ potassium chloride solutions at 298.15 K. *Journal of Chemical & Engineering Data*. 41(3):516-520

6.	González-Tello, P., F. Camacho, G. Blázquez, and F. J. Alarcón. 1996. Liquid− liquid equilibrium in the system poly (ethylene glycol)+ MgSO4+ H2O at 298 K. *Journal of Chemical & Engineering Data*. 41(6):1333-1336

7.	Perry, D., D. Momotenko, R. A. Lazenby, M. Kang, and P. R. Unwin. 2016. Characterization of nanopipettes. *Analytical chemistry*. 88(10):5523-5530
